# Chromatix: a differentiable, GPU-accelerated wave-optics library

**DOI:** 10.1101/2025.04.29.651152

**Authors:** Diptodip Deb, Gert-Jan Both, Eric Bezzam, Amit Kohli, Siqi Yang, Amey Chaware, Cédric Allier, Changjia Cai, Geneva Anderberg, M. Hossein Eybposh, Magdalena C. Schneider, Rainer Heintzmann, Fabrizio A. Rivera-Sanchez, Corey Simmerer, Guanghan Meng, Jovan Tormes-Vaquerano, SeungYun Han, Sibi Chakravarthy Shanmugavel, Teja Maruvada, Xi Yang, Yewon Kim, Benedict Diederich, Chulmin Joo, Laura Waller, Nicholas J. Durr, Nicolas C. Pégard, Patrick J. La Rivière, Roarke Horstmeyer, Shwetadwip Chowdhury, Srinivas C. Turaga

## Abstract

Modern microscopy methods incorporate computational modeling as an integral part of the imaging process, either to solve inverse problems or optimize the optical system design itself. These methods often depend on differentiable optics simulations, yet no standardized framework exists—forcing computational optics researchers to repeatedly and independently implement simulations with limited reusability and performance. These common problems limit the potential impact of computational optics as a field. Here we present Chromatix: an open-source, GPU-accelerated, differentiable wave optics simulation library. Chromatix builds on JAX to democratize fast, parallelized simulation of diverse optical systems and expand the design space in computational optics. Chromatix standardizes a growing collection of optical elements and propagation methods allowing a broad range of applications, which we demonstrate here for snapshot microscopy, holography, and phase retrieval. We demonstrate speed improvements of 2-6× on a single GPU and up to 22× on 8 GPUs.

## Introduction

Modern microscopy methods increasingly rely on computation as an integral part of the imaging process. This model-based approach to optics—integrating optical system design with algorithmic reconstruction or optimization—has had major implications for biological discovery. Single molecule localization microscopy can reveal individual protein complexes without requiring electron microscopy^1,2^ and 3D snapshot microscopy enables volumetric imaging at the frame rate of the camera, facilitating whole brain imaging at unprecedented temporal resolution^3–8^. For both techniques, using wave optics models to engineer point spread functions (PSFs) allows improved resolution in 3D^3,9–11^. Computer-generated holography uses wave optics models of light propagation to design precise optogenetic stimulation of many neurons simultaneously^12–15^. Integrating microscopy with wave optics models also allows measurement of optical properties that are otherwise difficult or slow to obtain, such as using diffraction tomography through a strongly scattering sample to obtain 3D refractive index distributions of transparent tissues from only intensity images, allowing high-contrast, label-free imaging of optically transparent model organisms like *C. elegans* or *D. rerio* ^16,17^. Increasingly, differentiable models of wave optics have also allowed for the end-to-end optimization of optical design and image processing algorithms for a variety of techniques such as quantitative phase imaging^18^, hyperspectral imaging^19^, extended depth-of-field^20^, monocular depth estimation^21,22^, localization microscopy^9^, lensless imaging^23^, and 3D snapshot microscopy^3^.

However, each computational optics method typically requires the researcher to program an optics simulation from scratch. These *de novo* simulations have differing conventions with other simulations, can be difficult to reuse in other applications, and are often computationally suboptimal due to the difficulty of programming fast optics simulations on modern computer hardware like graphics processing units (GPUs). Moreover, these simulations typically need to be differentiable to facilitate efficient optimization of the relevant optical parameters or to be easily combined with deep neural networks. A standard library providing differentiable wave optics simulations that make efficient use of GPUs is therefore desirable.

Here we describe Chromatix, a high-performance differentiable wave optics simulation library that could fill this gap in the field of computational optics. Drawing inspiration from modern deep learning frameworks like PyTorch^24^ and TensorFlow^25^ which construct neural networks from layers of mathematical operations, Chromatix enables researchers to describe optical systems as compositions of fundamental elements. This architectural parallel runs deeper than mere interface design: Chromatix shares the core requirements of differentiability for gradient-based optimization, scalability for tackling large-scale problems, and composability for rapid method development. Chromatix is built on JAX (Just After eXecution)^26^, a numerical computation library for Python that provides GPU acceleration and automatic differentiation (AD) to automatically and efficiently calculate gradients with respect to any input of differentiable functions. By leveraging JAX’s capabilities, Chromatix simulations can be seamlessly accelerated on GPUs and parallelized across multiple devices with minimal code changes (see Extended Data Figure 1), enabling easy integration with existing deep learning models, optimization loops, and even hardware control systems. Chromatix implements diverse optical models, from conventional lenses to diffractive elements like liquid-crystal-on-silicon (LCoS) spatial light modulators (SLMs), and various models of scalar and vectorial wave propagation through both free space and scattering media, enabling simulation of optical systems interacting with multiple-scattering (potentially anisotropic) biological samples whose amplitude and refractive index vary in 3D.

While there exist established tools for optical design such as Zemax^27^ or CODE V^28^ that do support wave optics, they are not efficiently implemented for the types of models we present here and are not differentiable or interoperable with deep learning models. Recently, a number of open-source differentiable optics simulation libraries have been released^29–32^; however, these libraries are either primarily ray-based^29,32^ (while the computational optics methods we are interested in are wave-based) or lack support for all the features that would be desirable for such a standard library in computational optics, e.g. multiple-scattering 3D samples or polarization. We provide a more detailed comparison of Chromatix versus other optics simulation software in Extended Data Table 1.

In this work, we demonstrate Chromatix’s ability to simulate and optimize various optical systems for biological imaging, showing simulations of widefield and snapshot fluorescence microscopes, phase contrast microscopes, and Fourier holography systems for optogenetics. In each case, we highlight Chromatix’s scalability, delivering results significantly faster than previous implementations—2-22× improvements through parallelization (depending on the problem)—while eliminating the challenges of correctly implementing wave optics simulations from scratch. We anticipate that this open-source library will democratize high-performance, scalable wave optics simulations, enabling exploration of a much richer design space in computational optics and accelerating innovation in biological imaging and beyond.

## Results

### Design and Implementation

The design of Chromatix is strongly inspired by modern deep learning frameworks. Here, we detail the principles that informed our design choices and implementation decisions. We argue that effective frameworks for computational optics, like those for deep learning, must embody three key characteristics: differentiability, composability, and scalability. We also discuss the high level implementation of Chromatix as it relates to these three characteristics.

#### Differentiability

Differentiability is the ability to calculate gradients, which can be used for gradient-based optimization (e.g., of the parameters of an optical simulation). For small, low-dimensional inputs, numerical differentiation can be sufficient, but for high-dimensional inputs (e.g., the pixels of an SLM) this becomes too computationally expensive to evaluate and it is preferable to use backpropagation of the gradients of each step of the simulation. When combined with the wide variety of possible optical models or neural networks, automatic differentiation (AD) becomes a desirable property. Common programming languages such as MATLAB^37^ and C do not provide general purpose AD, requiring gradients to be manually derived and implemented—an error-prone, time-consuming, and inflexible process. Modern deep learning frameworks^24–26^ provide AD: given a function, they can automatically calculate the gradient with respect to any parameter of that function as long as the function is differentiable with respect to that parameter. Similarly, Chromatix can automatically calculate the gradient with respect to any parameter of a simulation as it has been written using JAX.

Differentiability has already found several uses in optical design, enabling end-to-end design of computational optics systems for a variety of problems^3,9,18–23^. Differentiability also has the potential to improve solutions to inverse problems in optics. Traditional inverse problem approaches simplify both the sample and the optics^38–41^, whereas differentiable models can handle more realistic complexity. AD opens up a new class of gradient-based optimizers like Adam^42^, which can improve reconstruction fidelity by allowing arbitrarily complex physics simulations (e.g. scattering^16,36^ or sample deformation^43^) in the forward simulation. Importantly, these benefits come at nearly zero programmer effort with AD: once the forward model of the simulation has been defined, its gradients are automatically defined as well. Thus AD can also enable so-called self-calibrating algorithms^44^. As hardware has a finite accuracy, optimizing certain physical parameters (such as angle of illumination in tomography) together with the sample has been shown to improve fidelity^45,46^.

A different line of work replaces discrete voxel-based representations with neural network-based continuous representations, a concept known as implicit neural representations (INRs) or neural radiance fields (NeRFs)^43,47–49^. INRs have been applied to separation of motion artifacts from sample dynamics^43^, estimation of dynamic aberrations^48^, reconstruction of 3D quantitative phase of scattering samples^50^, and aberration correction without wavefront sensors or calibration measurements^47^. Here too, differentiable simulations are required to train these networks.

#### Composability

The principle underlying differentiability is composability: the gradient of a composition of two functions can be calculated from the gradient of each of those functions. Taking a somewhat broader and more practical view, we can interpret composability as being able to easily swap and replace components of a network, e.g. replacing the activation function in an MLP, without requiring changes to the rest of the system. This composability is possible due to standardization in the field of machine learning, which enables ML researchers to conveniently incorporate their colleagues’ advances by quickly replacing a function rather than having to rewrite their code from scratch. The field of optics stands in stark contrast: implementations are often project-specific, each with their own conventions and quirks, and a baseline to compare these codes to with respect to accuracy and speed does not exist. This practice is time-consuming, error-prone, and makes reproducing results challenging. Chromatix proposes a standard for wave-optics simulations to enable composition of a wide array of optical models, as shown in Figure 1. The experiments presented in this paper all share a common, well-tested codebase, and many more components are available in our documentation. We believe that the existence of both a standard library and baseline implementations can significantly speed up research and make it more reproducible.

**Fig. 1.**
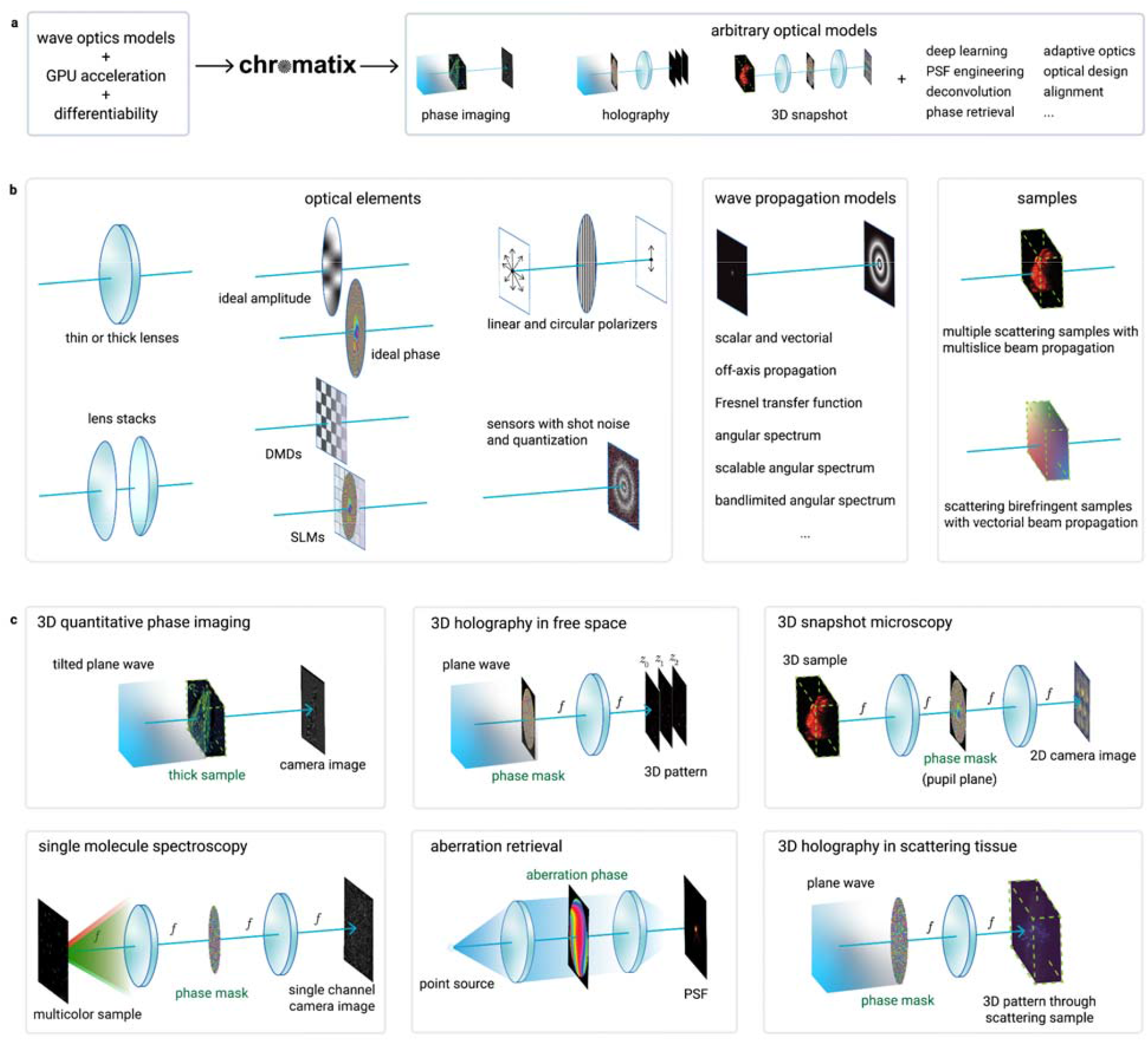
The design and components of Chromatix. **a**, Chromatix combines wave optics models, graphics processing unit (GPU) acceleration and differentiability in a single library, providing a unified modeling framework to allow a wide range of applications. **b**, Chromatix implements a wide range of optical elements such as lenses, sensors, free-space propagation models for scalar and vectorial waves ^33–35^, and complex scattering samples ^16,36^. **c**, These elements can be easily combined to simulate a wide variety of experimental systems and solve a wide range of problems in computational optics. Green highlighted elements indicate the element or sample that would be optimized in each application.

#### Scalability

Optics is moving to ever larger fields of view and higher resolutions, sometimes requiring large compute clusters for sample reconstruction. A key requirement for modern optics simulations is thus the ability to scale: researchers may want to run code on laptops for quick prototyping, but also easily scale up to GPU-clusters for large-scale sample reconstruction. Previous popular programming environments have made this difficult: NumPy runs only on central processing units (CPUs)^51^; MATLAB requires specific code for GPU usage and does not support general-purpose AD^37^; PyTorch^24^/TensorFlow^25^ make writing GPU code with AD relatively easy but it can be tricky to support multiple GPUs (both PyTorch and TensorFlow) or achieve good performance for typical operations in optical simulation which differ greatly from typical operations in neural networks (PyTorch). Writing device-specific programs for simulations requires significant effort and calcifies their capabilities, which is not appropriate for the fast iteration demanded by scientific research. Chromatix instead relies on JAX^26^ and its underlying XLA (Accelerated Linear Algebra) compiler to support fast optical simulation on CPUs, GPUs, and tensor processing units (TPUs) with only a single implementation (and without requiring custom lower level GPU code for fast operations as in PyTorch^24^). JAX also offers several functions to automatically vectorize code (i.e. parallelize a batch on a single GPU) or parallelize over multiple GPUs, independently of the description of the optical elements in an optical system^26^. For example, with only a couple of lines of changes to the code we can scale a 2D, single-wavelength simulation to a 3D, multi-wavelength simulation running on multiple GPUs in parallel (see Extended Data Figure 1 for code examples).

#### Implementation

In deep learning frameworks, these principles manifest themselves as models consisting of sequences of deep learning operations (layers). We observe a clear correspondence to optics, where optical systems consist of a sequence of optical elements and propagations. A key difference however is that the “hidden state” of an optical system has a clear physical meaning: it is the complex light field moving through the system. To completely describe this field, and thus the state of the system at any time, additional information such as the wavelength, polarization, and spatial sampling is required. The core idea behind Chromatix is that all this information can be encoded in a single, fundamental structure. Any optical element can then be written as a transformation of this structured field, and any optical system as a sequence of these elements. This allows Chromatix to model a wide variety of optical systems under a unified interface, which makes extending its capabilities straightforward.

### Experiments

We present six computational experiments demonstrating four major features of Chromatix: solving inverse problems to reconstruct samples, accelerating reconstruction and optical design using deep learning, composing modular optical elements and models in arbitrary ways, and scaling optical simulation speed by an order of magnitude. To do this, we showcase both reproductions of existing computational methods in optics that rely on wave models as well as *in silico* demonstrations of solving inverse problems in underexplored combinations of optical phenomena.

### Inverse problems for reconstructing samples

A microscope’s aberrations are usually assumed to be spatially invariant, due to the calibration and simulation complexity of simulating and measuring field-varying PSFs respectively. Kohli et al.^52,53^ argue that most imaging systems are rotationally equivariant due to the rotational symmetry of many optical elements, and that many common aberrations therefore vary only along the radius away from the center of the field of view (FOV), up to a rotation. Measuring only the variation of aberrations along this radial dependence requires substantially less calibration time and is much more efficient to model (linear versus quadratic scaling with the number of rows of the camera sensor). This efficiency enables tractable deconvolution of spatially-varying aberrations. Kohli et al. introduce “ring deconvolution microscopy,” which models a spatially-varying aberrated point spread function efficiently by exploiting the rotational invariance present in many standard microscopes using incoherently illuminated samples or fluorescent samples emitting incoherent light. We implemented this ring deconvolution method in Chromatix using the model shown in Figure 2b for UCLA (University of California Los Angeles) Miniscope^54,55^ data, a miniature widefield microscope (detailed in Methods). The microscope is modeled as a 4f-system with rotationally invariant Seidel aberrations in the Fourier plane. After estimating Seidel coefficients from calibration images^52^, the measured image of the sample is deconvolved using this rotationally invariant but spatially varying model of the PSF.

**Fig. 2.**
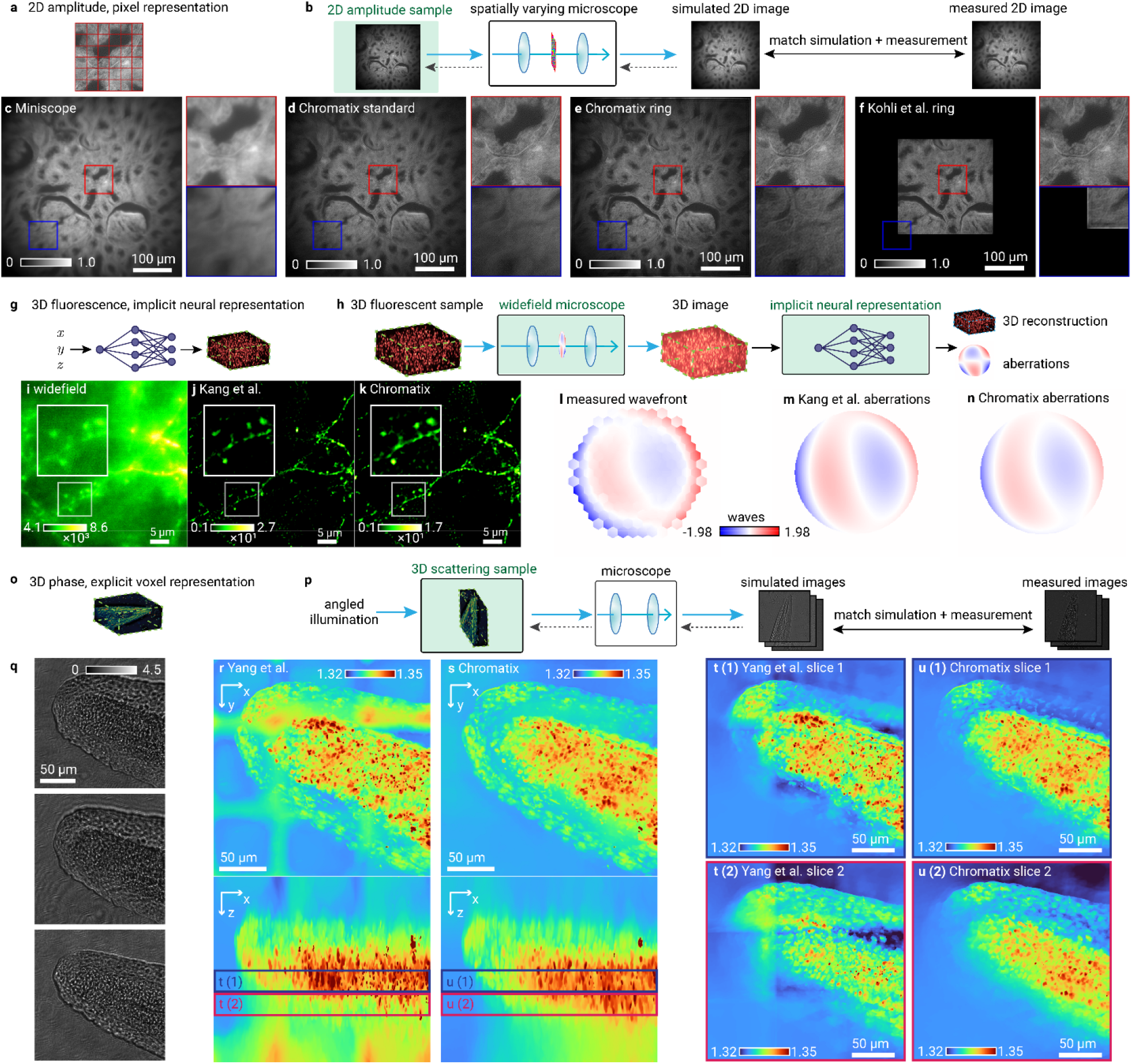
Chromatix solves inverse problems in multiple types of samples and sample representations. **a**, Sample type and representation for ring deconvolution microscopy. **b**, Implementation of ring deconvolution microscopy following Kohli et al., 2022. Highlighted regions in **c**-**f** show zoomed in cutouts from the center (red) and edge (blue) of the field of view on the bottom row. These images have been corrected to reduce vignetting for display purposes. Uncorrected images are shown in Extended Data Figure 2. Measured image of incoherently illuminated rabbit liver **c** from a Miniscope with a GRIN lens showing extreme spatially-varying aberrations across the field of view; **d**, Spatially invariant (standard) deconvolution demonstrating good quality in the center but degraded quality in the edges of the field of view; **e**, Chromatix rotationally invariant deconvolution demonstrating improved image quality across the entire field of view; and **f**, original PyTorch rotationally invariant deconvolution which does not parallelize and cannot fit the whole field of view on 1 H100 GPU (80 GB memory). **g**, Sample type and representation for computational aberration correction using implicit neural networks. **h**, COCOA self-supervised framework implementation using coordinate-based implicit neural networks for simultaneous aberration inference and 3D sample reconstruction from a single 3D measurement (Kang et al., 2024). **i-k**, Visual comparisons showing a maximum projection of a 4 µm slice from a raw aberrated measurement (**i**), original implementation reconstruction using PyTorch (**j**), and Chromatix reconstruction which better preserves the continuity of the dendrite (**k**). **l-n**, Validation of aberration recovery showing true measured wavefront using direct wavefront sensing (**l**), inferred aberration from original PyTorch implementation (**m**), and Chromatix inference of the aberration (**n**). **o**, Sample type and representation for 3D refractive index microscopy. **p**, Refractive index microscopy to recover the 3D refractive index distribution of a strongly scattering sample from intensity measurements. Raw measurements (**q**) from multiple angles of coherent illumination are used to reconstruct the full 3D refractive index map of the tail of a *D. rerio* embryo at 24 hpf, demonstrated by the original MATLAB reconstruction (Yang et al., 2024) of refractive index maximum projection **(r)** and the Chromatix reconstruction maximum projection **(s). t-u**, Slices through the full volume (colored borders denote slice regions) are shown to highlight the reduced grid artifacts in the Chromatix reconstruction (**u (1), u (2)**) compared to the original MATLAB reconstruction (**t (1), t (2)**). For all panels, green highlights denote optimized parameters and dashed gray arrows denote propagation of gradients during iterative optimization.

We show respectively the measured image of incoherently illuminated rabbit liver, ring deconvolution using Chromatix which recovers detail across the whole FOV, standard deconvolution which only recovers detail in the center of the FOV, and ring deconvolution from the original implementation by Kohli et al. (Figures 2c-f, Extended Data Figure 2). Note that, compared to the original implementation, our reconstruction has a significantly larger FOV; our implementation was able to reconstruct a larger image by parallelizing across multiple GPUs. The original implementation fails to reconstruct the entire field of view of the camera (without suffering significant degradation in reconstruction speed) due to overflowing the memory limitations of a single GPU (48 GB for an RTX 8000 or 80 GB for an H100). Chromatix’s implementation is also significantly faster, showing a 4.5× speedup versus the original PyTorch implementation on a single GPU and scaling up to almost 19× when using 8 GPUs as shown in Figure 5.

In addition to voxel grids, Chromatix also enables implicit neural representations (INRs) that in some cases improve the optimization loss landscape^48^. CoCoA (Coordinate-based neural representations for Computational Adaptive optics)^47^ jointly reconstructs the sample (represented as an INR) and the aberrations (represented as Zernike coefficients) in a self-supervised fashion, as shown by the Chromatix implementation of the COCOA method in Figure 2g-h. Contrary to the previous section, the aberrations here are modeled as spatially-invariant and the sample emits incoherent fluorescent light, rather than transmitting incoherent illumination. We show in Figures 2i-k respectively a maximum intensity projection of a 4 µm slice through a measured widefield mouse neuron volume, the reconstruction of the unaberrated sample from the original implementation, and finally the reconstruction using Chromatix. We note that the Chromatix implementation retains more uniform dendrites which become spotted in the original implementation. Chromatix is also twice as fast at performing the reconstruction when using a single GPU, or almost 9× faster when using 8 GPUs (Figure 5). Further, on a fluorescent bead dataset with controlled, intentional aberrations provided by Kang et al., the Chromatix implementation recovers the applied Zernike mode coefficients with a root-mean-square (RMS) of 3.56 nm versus an RMS of 6.97 nm for the original implementation^47^ (RMS is computed on the three non-zero Zernike modes that were used to intentionally aberrate the system; see Extended Data Figure 3).

The loss of detail in the original implementation may be mitigated by increasing the number of layers of the INR as reported by Kang et al., but here we compare reconstructions using identical network architectures. The Chromatix implementation uses a paraxial approximation for the field at the pupil plane while the original implementation by Kang et al. mixes the exact model for the field at the pupil (requiring a higher sampling rate to avoid aliasing the propagation kernel) with a paraxial approximation of the second lens of the 4f system. In this case, the fully paraxial approximation used by Chromatix results in an improved reconstruction of the dendrite as shown by Figure 2i-k. This demonstration highlights not only our increased performance but also the utility provided by a standard set of models like Chromatix when faced with the potential for model mismatch.

Chowdhury et al. ^16^ show that computational imaging is also able to quantitatively recover the 3D refractive index distribution of strongly scattering samples (i.e. beyond the first Born approximation) from intensity measurements, a quantity otherwise inaccessible to conventional widefield light microscopy. The sample (the tail of a *D. rerio* embryo 24 hpf) is coherently illuminated at different angles according to the setup described by Yang et al. ^17^, and the refractive index (Figure 2r) is recovered by matching the measurements (Figure 2q) to a differentiable simulation (Figure 2p) of imaging through the scattering sample. Typical samples are beyond the single-scattering regime and are hence modeled using a multislice approach ^56^. While we use exactly the same forward model as the original MATLAB implementation ^17^, our implementation is 3-13× faster than the original implementation on equivalent volumes (bringing reconstruction time from hours to minutes). Our implementation is also far fewer lines of code (approximately 25 lines for a differentiable simulation in Chromatix versus approximately 107 lines in the original implementation^16,17^) and more flexible compared to the original implementation with respect to changes in the forward model or optimization parameters due to the use of automatic differentiation. This increase in speed allows us to choose more appropriate reconstruction settings so that the large grid artifacts in the original reconstruction (Figure 2r) are removed in the Chromatix reconstruction (Figure 2s).

### Programmable optics and deep learning

The recent commercial availability of SLMs has made fine-grained control of light possible through millions of controllable pixels. These degrees of freedom are often used for holography, but can also be used to design a point spread function for a specific purpose^57^ such as volumetric snapshot imaging for fluorescent 3D samples. The dimensionality of this optimization problem requires gradient-based optimization to be effectively solved and is well-suited to GPU acceleration. Deb et al.^3^ introduce a deep learning method for engineering 3D snapshot PSFs that combines a programmable microscope design (the Holoscope) along with a neural network to reconstruct fluorescent 3D volumes from 2D snapshot images taken by the microscope (Figure 3a). The parameters of the neural network along with the pixels of the programmable phase mask implemented by an SLM are jointly optimized using a differentiable simulation of the microscope. In this snapshot microscope, the PSF essentially acts as a compression function from the 3D volume to the 2D image. The volume is then reconstructed from this image using a FourierNet neural network, so that structural priors of the sample type can be taken advantage of by both the optics and the computational reconstruction algorithm.

**Fig. 3.**
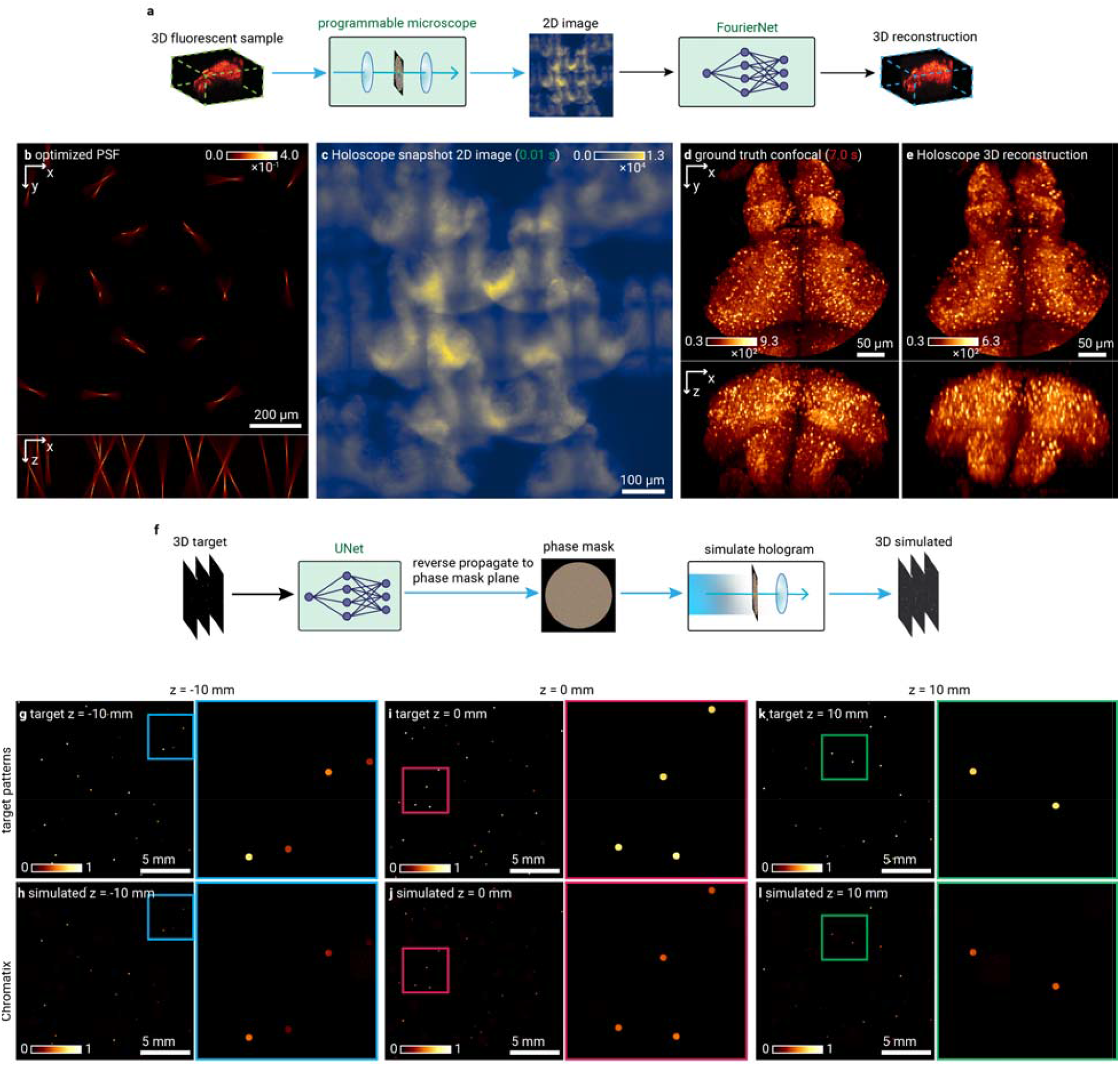
Chromatix enables deep learning for optical design and acceleration of inverse problems. **a**, Holoscope3 implementation in Chromatix showing a programmable 3D snapshot microscope, compressing volumetric information into a 2D representation with subsequent FourierNet reconstruction. **b-e**, Holoscope demonstration using Chromatix showing the sample-specific point spread function (**b**), simulated 2D image of the simulated 3D sample (**c**) (approximately 0.01 seconds to capture), ground truth 3D volume (**d**) (approximately 7.0 seconds to capture via confocal microscopy assuming a scan speed of 100 ns per voxel), and Chromatix-enabled 3D reconstruction (**e**) from the single simulated 2D image. **f**, DeepCGH architecture utilizing a UNet and a propagation step to directly generate a hologram in a single feedforward step from target 3D patterns. **g-l**, Demonstration of DeepCGH12 using Chromatix showing requested stimulation patterns at three planes spaced 10 mm apart around the focal plane (**g, i, k**) and their resulting simulated intensity distributions (**h, j, l**). Colored insets show detail of the 3D patterns at each plane. Intensity values in **g**-**l** are normalized. For all panels, green highlights denote optimized parameters of either neural networks or optical systems.

This microscope design can therefore be programmed to function as a snapshot microscope optimized for various sample types, while using exactly the same hardware. The microscope is modeled using a 4f system with an SLM (phase mask) in the Fourier plane and is optimized for whole-brain imaging of fluorescently labeled *D. rerio* larvae. The PSF of the 4f system is simulated with coherent propagation, and the image is simulated as the incoherent sum of these PSFs which is efficiently implemented as a convolution of the PSF and the sample intensity. We show the learned PSF (Figure 3b), the simulated 2D measurement of a virtual zebrafish volume (Figure 3c), and Figures 3d and 3e the ground truth volume and simulated reconstruction^3^. Chromatix reproduces the original results of Deb et al.^3^ nearly exactly: on a test set of 10 volumes and their simulated images, reconstruction networks trained with identical PSFs offer a structure similarity index measure (SSIM) of 0.979 ± 0.003 (higher is better) for both Chromatix and the original implementation^3^ (not significantly different, see Extended Data Figure 4). Chromatix also outperforms the original implementation^3^ in training speed by approximately a factor of 7× (Figure 5). Practically, this reduces the optimization time for a single PSF from weeks to days.

Spatial light modulators also enable computer generated holography systems for optogenetics, where 3D holographic stimulation patterns are used to perturb neural activity in the brain. Most holography systems rely on some form of iterative optimization (e.g. Gerchberg-Saxton^38,39,58,59^) to find the phase to display on the SLM. While this produces accurate solutions, iterating does become problematic when speed is paramount. For optogenetics, point cloud holography can be used to stimulate multiple neurons without iterative optimization of phase patterns, but this only allows for placing copies of a single pattern at the desired locations^15^. Due to the interest in holography for displays, fast holography algorithms for arbitrary patterns have emerged that use neural networks to quickly generate a hologram given a target pattern^46,60^. Applied to optogenetics, DeepCGH^12^ also demonstrates fast computer-generation of holograms by training a neural network to generate phase patterns from intensity images of arbitrary 3D patterns in a single feedforward inference step. We implemented DeepCGH^12^ as shown in Figure 3f. We show the desired target patterns and resulting simulated pattern at three different depth planes using the phase pattern produced by the DeepCGH method in Chromatix (Figure 3g-l). We achieve nearly identical results as the original TensorFlow implementation: on a test set of 16 target patterns, Chromatix achieves an SSIM (higher is better) of 0.985 ± 0.001 versus 0.982 ± 0.001 for the original implementation (not significantly different, see Extended Data Figure 5). Our implementation is approximately 17 lines of code for a differentiable hologram simulation versus 33 lines in the original work^12^. While achieving the same quality, Chromatix provides a 2.5× performance improvement on a single GPU, which increases to over 10× when using 8 GPUs in parallel (Figure 5).

### Flexible modeling with optical building blocks

Because Chromatix models are constructed from components that can be flexibly combined as shown in Figure 1, we can straightforwardly construct complex optical models and also optimize them with arbitrary objective functions. We show another programmable microscope modeled as a 4f system with an SLM in the Fourier plane, followed by a neural network-based reconstruction step (Figure 4a-f). The objective in this demonstration is to optimize the PSF of this programmable microscope to perform spectroscopic single-molecule localization^61^ from a single snapshot image, i.e. to reconstruct multicolor point sources using only a single-channel image. The simulated samples consist of several point sources incoherently emitting fluorescence at 25 wavelengths from 400 to 650 nm which are simulated in parallel using Chromatix. We train the neural network to reconstruct the multicolor sample at the corresponding 10 nm intervals, giving us a hyperspectral cube from a single-channel 2D measurement. As shown in Figure 4c, the optimized PSF allows visual classification of the color of these point sources on a monochrome simulated camera image (Figure 4d) by taking advantage of different fringe patterns for different wavelengths. The reconstruction (Figure 4f) reasonably matches the true colors of the points in the sample (Figure 4e). We highlight that here we are optimizing the same programmable microscope model that was used for snapshot microscopy in Figure 3a, but for an entirely new combination of sample type and objective.

**Fig. 4.**
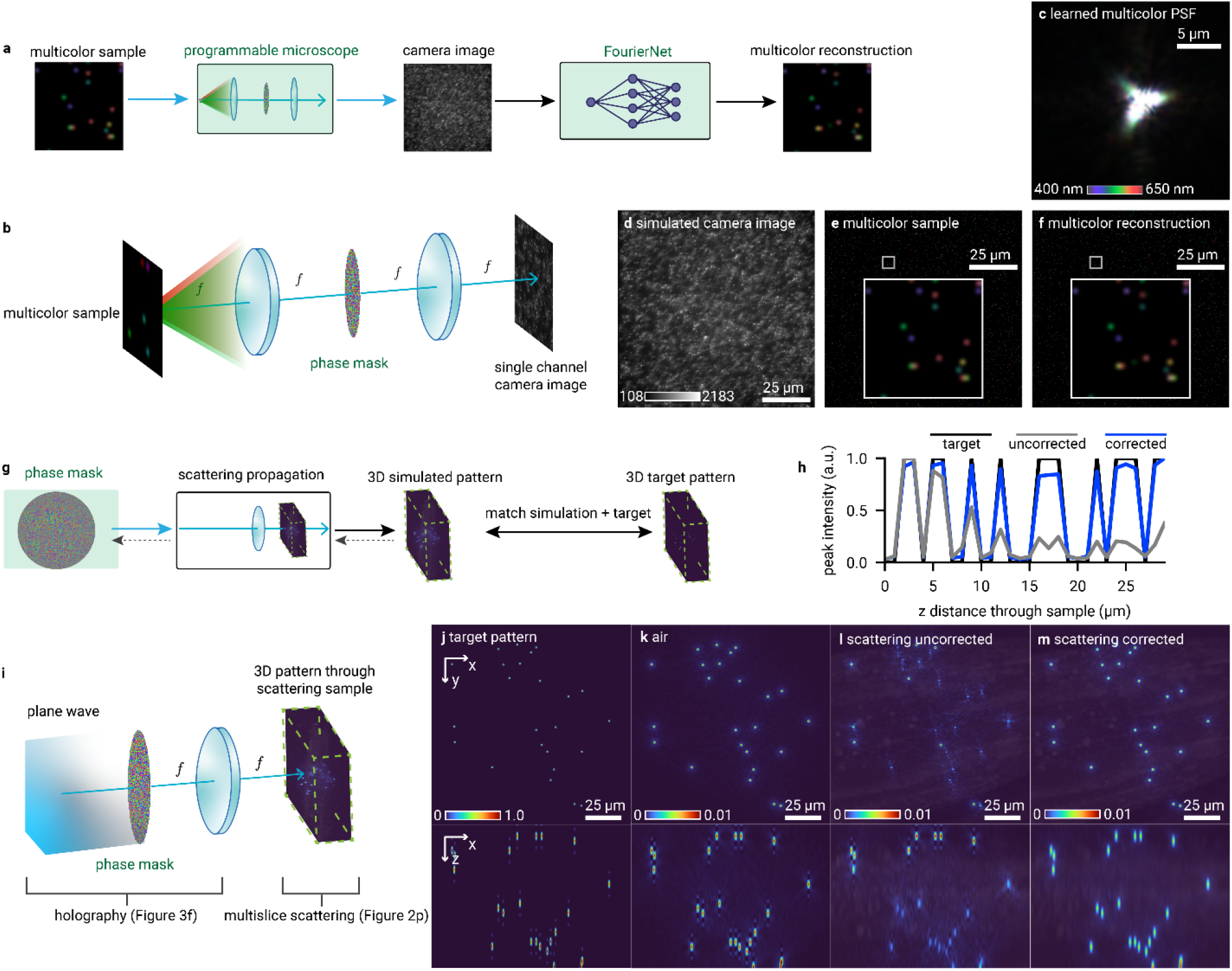
Chromatix enables arbitrary combinations of optical models. **a**, Demonstration of PSF engineering for spectroscopic single-molecule localization microscopy using deep learning, where a neural network reconstructs both the structure and spectrum of a sparse 2D point sample from a single-channel image. **b**, Microscope model for multicolor PSF optimization with a single spatial light modulator (SLM) in the Fourier plane. **c**, Optimized multicolor PSF for spectral imaging using a neural network, with different wavelengths/colors overlaid. **d**, Simulated single-channel image of multicolor fluorescent point sources. **e**, True simulated multicolor point sources, with different wavelengths/colors overlaid. **f**, Reconstructed multicolor point sources, with different wavelengths/colors overlaid. **g**, Iterative optimization workflow generating optimal phase masks for scattering-compensated holography. **i**, Chromatix model of holographic pattern formation through scattering media which combines the holography model of Figure 3f and the scattering sample model of Figure 2p. **h**, Axial intensity distribution (normalized within the range 0-1) showing scattering-induced non-uniformity of the stimulation pattern along the direction of propagation, which can be corrected through optimization. **j-m**, Visual evolution of holographic pattern quality. Target pattern (**j**) is well matched by an optimized hologram simulated through free space (**k**) but not well matched when simulated with scattering through a sample with a known 3D refractive index distribution (**l**), unless we use Chromatix to correct the hologram through the scattering volume (**m**). For all panels, green highlights denote optimized parameters and dashed gray arrows denote propagation of gradients during iterative optimization.

Arbitrary combinations of wave optics models can also open up further applications of differentiable simulations to biological research. Optogenetic experiments rely on complex, 3D patterns of light to selectively control the behavior of neurons, often using holography^12,14,15^. 3D holography in itself is challenging, but this is compounded in optogenetics by the scattering nature of the biological tissue. As the light propagates through the tissue, it gets scattered, possibly activating the wrong neurons and increasing phototoxicity ^62–64^. In Figure 4g-m we show how Chromatix can be used to optimize the desired holographic pattern in such a strongly scattering tissue by imaging the scattered pattern. We model this scenario as a plane wave incident on a phase mask (SLM), focusing through a thin lens, and finally propagating through a scattering volume. The propagation through the volume is modeled using the multislice beam propagation (MSBP) method (using the same code to model the scattering sample as Figure 2p), and we observe in simulation the intensity throughout the entire volume. Without correction, the stimulation is uneven due to the unaccounted-for scattering, as shown in Figure 4h. Once we include the observed scattered intensity as feedback in the optimization, we obtain a near uniform stimulation (blue line in Figure 4h) across the whole axial range. This serves as an *in silico* demonstration that Chromatix enables researchers to rapidly implement and iterate on novel ideas, transforming intuition into tangible results.

### High performance through parallelization

To demonstrate the computational performance and scalability of Chromatix, we benchmark iteration speed for all of training and optimization problems we have presented. Chromatix has superior performance across all reproduced optical methods, ranging from 2-6× on single GPUs to 22× faster on 8 GPUs in the best case as shown in Figure 5. Single GPU performance improvements are typically due to less overhead after compilation through JAX as compared to other implementations in MATLAB, PyTorch, or TensorFlow. In general, order of magnitude improvements are possible via parallelization with Chromatix. Importantly, these parallelization schemes can be implemented on Chromatix models with virtually no changes to the code defining the models due to Chromatix being implemented in JAX ^26^. This acceleration allows Chromatix to scale to solving large problems, and also makes existing inverse problem solutions significantly more tractable as demonstrated by the larger field of view for the Chromatix reconstruction in ring deconvolution microscopy (Figure 2a-f) as well as the dramatic decrease in optimization time for refractive index microscopy (Figure 2o-u) and snapshot PSF optimization with deep learning (Figure 3a-e).

**Fig. 5.**
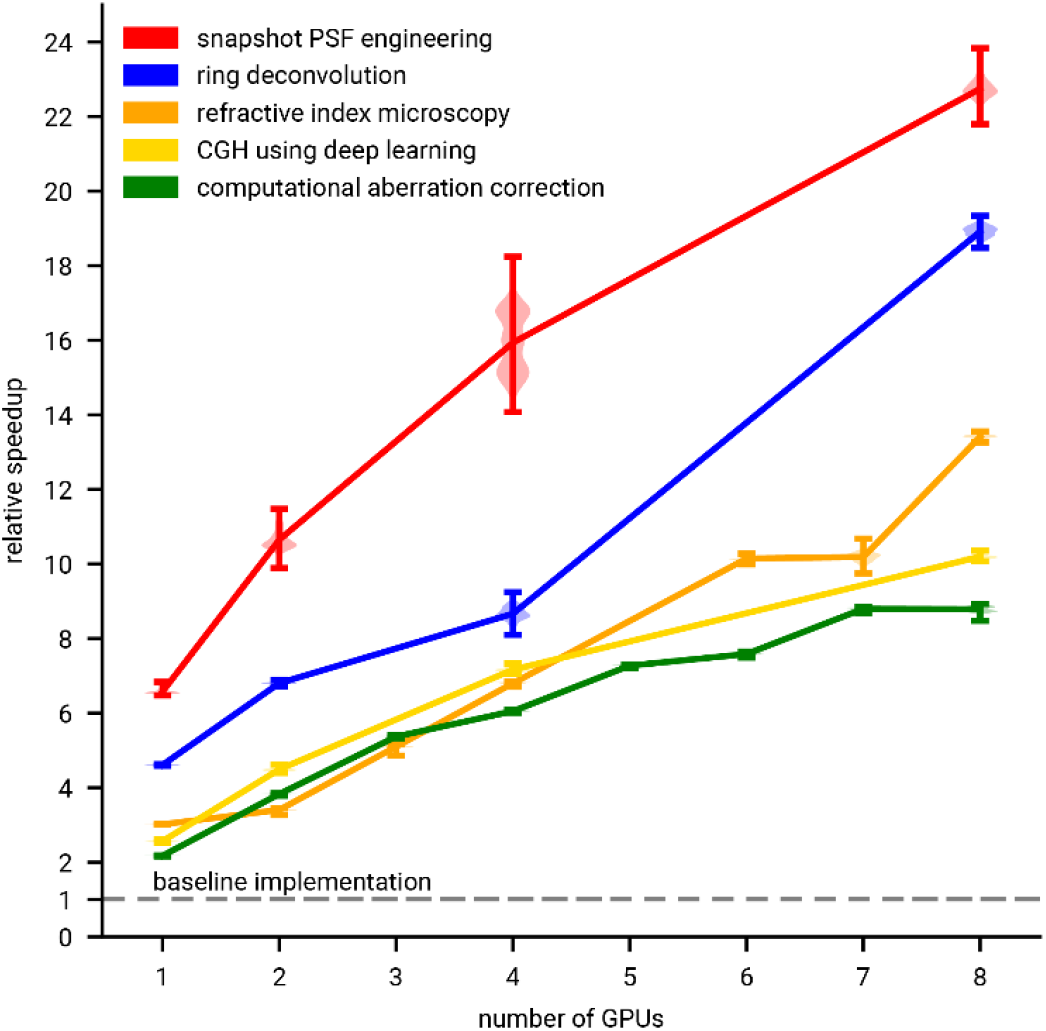
Chromatix is the fastest implementation of existing computational optics methods. Vertical axis shows relative speedup of Chromatix on 1 - 8 GPUs compared to the original implementations of each method as the baseline (represented via the gray dashed line at 1×). Points are centered on the mean relative speedup for each method. Error bars show standard error of speedups on individual iterations of the optimization for each method. The distribution for each method is visualized as a violin plot in a lighter shade. See Methods for details. For all methods, speedups are relative to the original single GPU implementations.

## Discussion

We introduced Chromatix, a differentiable wave optics simulation library that standardizes a wide and growing range of wave optics models. While many computational optics techniques incorporate wave optics models, including deconvolution, PSF engineering, holography, and phase retrieval, these models must typically be programmed from scratch for every project with differing mathematical conventions and limited reusability. These de novo implementations are often not maximally performant or easily parallelizable (Figure 5), which can also reduce the quality of results obtained by these methods due to the resulting practical limitations in reconstruction size or optimization settings as we demonstrate in Figure 2. A standard library for wave optics that is reasonably performant while remaining easy-to-use for research could address these problems in the field of computational optics.

By recreating a set of models and methods spanning all aforementioned techniques (Figures 2 and 3), we demonstrated that Chromatix provides a rich modeling framework to become the standard library in the field. The collection of models in Chromatix spans scalar and vectorial wave propagation, complex multiple-scattering 3D samples, and a variety of optical elements including lenses, SLMs, and sensors. In addition to this computationally efficient and standardized collection of optical elements, Chromatix models are automatically differentiable which enables integration with neural networks. These networks can be used to represent samples as demonstrated in Figure 2g-n (computational aberration correction^47^) for either memory-efficient representations^49^ or improved image reconstruction quality^43^. They can also be used to perform end-to-end joint optimization of optical systems and the neural networks solving the corresponding inverse problems as we demonstrated in Figure 3 (3D snapshot PSF engineering^3^ and accelerated holography^12^) and Figure 4a-f (spectroscopic single-molecule localization PSF engineering^61^).

Several of the major features in Chromatix were added by participants at hackathons we held to teach Chromatix. Across three hackathons at both HHMI Janelia Research Campus and Johns Hopkins University, a total of 32 participants with backgrounds in biology and optics—but not necessarily advanced numerical computing—were able to implement models and features required for their research in just a few days. Participants at subsequent hackathons and other researchers could then build on the work added to the library due to the common framework imposed by Chromatix. This highlights that Chromatix’s accessibility allows researchers to explore state-of-the-art computational optics, and also that standardization of models makes it easier to share and disseminate their work. We expect this trend of open-source contributions to continue, adding an ever-increasing collection of models which would be necessary for use as a standard library in computational optics.

We are continuing to develop Chromatix in several ways to address its limitations. Chromatix models some optical elements such as lenses in idealized ways (with experimental support for thicker lenses); the library of optical elements can be expanded with more realistic modeling of existing lens geometries with lens coatings^27^, and sensors with noise patterns and pixel responses which are currently missing from Chromatix. Further, while Chromatix focuses on wave-based modeling of optics, it may be necessary to use ray-based modeling for computationally efficient simulation of complicated series of lenses or other surfaces. Yang et al.^65^ show how to combine ray-based models with diffractive models, providing a path for Chromatix to incorporate ray-based models of lenses in the future. Computationally efficient modeling of forward and backward scattering through inhomogeneous samples can be implemented via the modified Born series (MBS)^66^. Chromatix also does not support rigorous coupled-wave analysis, which is often used to efficiently simulate and design metasurfaces^67^. Hazineh et al.^68^ show an efficient differentiable method for this inverse problem that could allow Chromatix to support metasurface design. Finally, Chromatix currently models completely coherent propagation, which can be expanded to include support for partially coherent propagation models^69^ as well as changes in the spectrum of light due to propagation^69^.

Looking further, we envision a wide range of possible applications for fast simulations provided by Chromatix, such as architecture search for designing optical systems (analogous to neural architecture search for optimizing network architectures^70^) or integration with hardware control software like Micro-Manager^71^/Pycro-Manager^72^ to enable automatic calibration or alignment with hardware-in-the-loop techniques^46^ (increasing reproducibility for computational optics techniques); both of these applications would benefit from being able to evaluate many simulations quickly and in parallel. Just as accessible deep learning libraries enabled deep learning researchers to explore a rich space of neural network architectures and methods, we expect the standardization of computationally efficient and scalable wave optics models in Chromatix to enable the exploration of a rich space of computational optics methods that may have previously been impractical to implement.

## Acknowledgements

The authors would like to thank Hari Shroff and Xuesong Li for their careful reading of the manuscript.

## Funding

This work was funded by the Howard Hughes Medical Institute (S.C.T., D.D., G.J., C.A., M.S.). B.D. was supported by a Grant from the German-Israeli Foundation for Scientific Research and Development (GIF, Grant number G-1566-413.13/2023). C.J. and lab (Y.K.) are supported by the National Research Foundation of Korea (NRF) grant funded by the Korea government (MSIT) (No.2023R1A2C3004040) as well as The Ministry of Trade, Industry and Energy (MOTIE) and Korea Institute for Advancement of Technology (KIAT) through the “International Cooperative R&D program (Task No. P0028516)”. L.W. is a Chan Zuckerberg Biohub SF investigator; U.S. Air Force Office of Scientific Research award no. FA955-22-1-0521. S.C. and lab (S.Y. and S.C.S.) are supported by the Chan Zuckerberg Initiative (2021-225666, 2023-321173), the National Institutes of Health (NIBIB R21□EB033629, NIGMS R35□GM155424), and the University of Texas at Austin; Cockrell School of Engineering; Chandra Family Department of Electrical and Computer Engineering. S.H. is supported partly by grant 2021-234544 from the Chan Zuckerberg Initiative DAF, an advised fund of the Silicon Valley Community Foundation. N.C.P. and lab (C.C., M.H.E., and J.V.T.) are supported by a Beckman Investigator grant from the Arnold and Mabel Beckman Foundation, a Sloan Fellowship in Neurosciences from the Alfred P. Sloan Foundation, and a Kavli Innovation Grant.

## Author Contributions Statement

All authors contributed to the development of the Chromatix library. D.D. and G.J. contributed equally by leading the development of the library and conducting the computational experiments for the manuscript. D.D., G.J., and S.C.T. wrote the manuscript with contributions from all authors.

## Competing Interests Statement

We declare that none of the authors have competing financial or non-financial interests as defined by Nature Portfolio.

## Methods

### Ring deconvolution microscopy (Figure 2a-f)

#### Visualized results

All procedures and data follow the original work by Kohli et al.^52^ We model a microscope with spatially varying aberrations, namely the UCLA Miniscope v3^54^ with Ximea MU9PM-MBRD 12 bit, 2.2 micron pixel sensor^52^. We show deconvolution of rabbit liver tissue embedded on a glass slide illuminated with 550 nm incoherent light, also shown originally in Kohli et al.^52^ Imaging is described as the incoherent sum of coherently simulated PSFs with spatially varying aberrations. Ring convolution efficiently models spatially varying aberrations by convolving each ring of a sample (the pixels at a certain radius from the center of the FOV) with the aberrated PSF corresponding to that ring^52^. Because Seidel aberrations are rotationally invariant, the PSF can be simulated at each pixel along a single diagonal path away from the center of the FOV, which defines up to a rotation the shape of the PSF at any other pixel^52^. While the original implementation in Kohli et al.^52^ using PyTorch could only reconstruct an image of size 1024 × 1024 pixels due to memory limitations on a single GPU, the Chromatix reconstruction could process the full camera image of 2048 × 2048 pixels while modeling spatially varying aberrations by parallelizing the reconstruction across multiple GPUs.

For reconstruction, we normalize pupil coordinates (u, v) across the camera image to have magnitude less than or equal to 1 within the original cropped 1024 × 1024 pixel region used by Kohli et al.^52^, and scale coordinate magnitudes beyond 1 outside of this region. This allows us to use the original estimated Seidel coefficients from the calibration performed by Kohli et al.^52^ for the Miniscope data: 0.85, 0.56, 0.25, 0.29, and 0 waves for the five Seidel aberrations. The spatially varying PSF is simulated at all pixel coordinates along one radial line from the center of the image to one corner, and these PSFs are rotationally Fourier transformed for use in ring convolution, as outlined in Kohli et al.^52^ The forward model can then be defined by the ring convolution between the spatially varying PSFs and the sample. Deconvolution is performed by initializing the sample as the measured camera image, and then iteratively performing gradient descent with the Adam optimizer ^42^ to optimize the pixels of the reconstruction by comparing the loss between the simulated image from the forward model and the measured camera image^52^.

#### Scalability results

We demonstrate reconstructions parallelized across 1, 2, 4, or 8 GPUs (NVIDIA H100s). Parallelization is performed by spreading different rings (different radii) of the ring convolution across different GPUs. To obtain the final image, the different rings are synchronized across the different GPUs after the convolution is performed. We reconstruct a 1024 × 1024 pixel image so that the original implementation (which is single GPU only) can complete a reconstruction. We reconstruct a simulated image of point sources rather than an experimental image, because we are only concerned with measuring the reconstruction iteration speed and not reconstruction quality. We measure the iteration time to perform the forward simulation and gradient descent update of the reconstruction with the Adam optimizer.

### Computational aberration correction using coordinate-based neural networks (Figure 2g-n)

#### Visualized results

All procedures and data follow the original work by Kang et al.^47^. We model a widefield microscope with Zernike aberrations from ANSI (American National Standards Institute) indices 3-14 (without normalization). The sample shown is a 3D volume of fixed mouse brain slices (Thy1-GFP line M) with a number 1.5 cover glass (0.16 to 0.19□mm thickness) tilted at 3° on top to emulate a glass cranial window ^47^. The sample is excited in widefield by an expanded 488 nm laser beam. The emitted fluorescence was collected by a 1.1 NA objective lens, relayed to the camera by a 4f system. We recover a similar corrected (deconvolved) sample and estimated aberrations of the detection path as the original implementation in PyTorch.

We first optimize an INR to recreate the measured widefield volume by minimizing the structured similarity loss between the output of the network and the measured widefield volume. The implicit neural representation used to represent the volume takes as input an x-y location (radially encoded) and outputs the pixel intensity values for that location across all 200 z-planes of the sample. We then perform a second stage of optimization of the implicit neural representation using the forward model of the aberrated microscope. Here, we simulate imaging the volume output by the implicit neural representation using the forward model and minimize the structured similarity loss between the simulated widefield volume and the measured widefield volume. The simulation of the microscope requires first simulating the aberrated coherent PSF of the system and then convolving the 3D PSF with the 3D sample volume. In this second stage, we also update the Zernike coefficients of the forward model of the aberrated microscope. The final output is an implicit neural representation of the deconvolved sample and the Zernike coefficients representing the aberration of the microscope.

#### Scalability results

We measure iteration time during the second stage of training, including the time to perform a forward pass of the neural representation, a forward simulation of the aberrated microscope, and a backward pass to update the neural representation parameters as well as the Zernike coefficients of the aberrated microscope model using the Adam optimizer. We simulate a measured volume of point sources rather than the experimental data used in the visualization results to demonstrate the speed of optimization of Chromatix across 1 - 8 GPUs (NVIDIA H100s).

### Refractive index microscopy (Figure 2o-u)

#### Visualized results

All procedures and data follow the original work by Yang et al.^17^. We show a reconstruction from the tail of a zebrafish (*Danio rerio*) embryo (24 hpf). Images are measured by illuminating the transparent sample with coherent light from a 660 nm red laser (Thorlabs, S4FC660)^17^. This light passes through the sample at a specific angle, and then scatters through the transparent sample^17^. The light scattered through the sample at the focal plane of the objective (0.37 NA) is measured by the camera^17^. Many images are collected with different angles of illumination. Measured images are downsampled via cubic interpolation to 80% of their original size in both height and width, as performed by Yang et al. Certain images are manually omitted from use in the iterative reconstruction process for each field of view, as outlined in the data and reconstruction settings provided by Yang et al. Using the measured images, we reconstruct a volume of size 400 × 800 × 800 voxels (FOV 1) at a voxel size of (0.43125 µm z, 0.2875 µm y, 0.2875 µm x).

Reconstruction is an iterative optimization process over the voxels of the sample. Each voxel represents the change in refractive index at that location in the sample. The reconstruction is initialized to an all zero change in refractive index^17^. Images are simulated from each angle of illumination that was measured using multislice propagation through the scattering reconstruction, and the mean squared error between each simulated image and each measured image is computed^17^. The gradient of this loss is used to update the reconstruction via gradient descent. Once all images have been used to update the reconstruction, the current reconstruction is smoothed via a proximal total variation operator to a relative error of 10^-4^. This smoothed reconstruction is updated from the previous smoothed reconstruction using Nesterov accelerated gradient^17^. This whole process (forward simulating and updating from all the images) is repeated until the reconstruction converges^17^.

#### Scalability results

We reconstruct a volume of 400 × 800 × 800 from FOV 1 where we use all 168 images. We measure the iteration time to perform one forward simulation and backwards pass to compute the gradient descent update, not including any regularization. We split these 168 images across either 1, 2, 3, 4, 6, 7, or 8 GPUs (NVIDIA H100s). For a given number of GPUs, we forward simulate and compute gradients for that number of images in parallel. The gradients from this batch of images are averaged across all GPUs to compute the update for the given set of images. We report iteration time as the time to process a single image by dividing the measured iteration time for a batch by the size (number of images) of the batch.

### Snapshot PSF engineering using end-to-end deep learning (Figure 3a-e)

#### Visualized results

All procedures and data follow the original work by Deb et al. We follow the two-stage optimization procedure outlined in Deb et al., where we first optimize a weak FourierNet reconstruction network and phase mask to obtain a snapshot PSF for zebrafish larvae (*Danio rerio*) at 6-8 dpf ^3^. We then optimize a larger FourierNet reconstruction network while fixing the phase mask to obtain higher quality reconstructions. This optimization process uses 58 volumes of pan-neuronally labeled larval zebrafish (expressing nuclear-restricted GCaMP6s/7f) that were scanned at high resolution using a high resolution confocal microscope. These volumes were all resampled to the same pixel size (1.0 µm z, 1.5 µm y, 1.5 µm x) for training. The simulated snapshot microscope design images a 386 µm diameter region of the whole zebrafish that is 250 µm tall, which simulates imaging zebrafish in 3D with an aperture in the system blocking light from the sample outside of the desired range. The simulated camera field of view is 1132.8 µm.

Optimizing a PSF involves sampling different 386 µm diameter regions from the 58 total training volumes of larval zebrafish, applying affine transformation and brightness augmentations, simulating imaging the fluorescent volume at a wavelength of 513 nm using the snapshot microscope, and then reconstructing the 3D image from the 2D simulated image. Simulating imaging in this stage of optimization means first simulating the PSF, and then convolving each plane of the PSF with each corresponding plane of the sample after downsampling the PSF plane to the resolution of the sample plane. The final 2D image is the sum of these planewise convolutions. We then jointly update both the weak neural network parameters for reconstruction and the phase mask parameters (pixels) using a loss that averages the mean squared error between the reconstruction and the true volume as well as the mean squared error between the high pass filtered reconstruction and high pass filtered true volume. In the second stage of training, we use a more powerful neural network and fix the phase mask parameters.

#### Scalability results

We report iteration times for the first stage of training, where we are optimizing a phase mask (2560 × 2560 pixels) by jointly optimizing phase mask pixels and reconstruction neural network parameters. Simulating the PSF at 64 planes spanning 250 µm where each plane of the PSF has 2560 × 2560 pixels means we simulate a PSF containing 0.419 unique gigavoxels. The neural network reconstructs 64 planes spanning 250 µm where each plane of the volume has 512 × 512 pixels. Here, for both the original implementation by Deb et al. using PyTorch and the Chromatix implementation, we report iteration times across 1, 2, 4 or 8 GPUs (compared on NVIDIA RTX 8000s due to limitations of old PyTorch code). Parallelization for both the original implementation and the Chromatix implementation involves splitting the simulation of different planes (both for the PSF and for the imaging) across different GPUs, so that each GPU simulates its own chunk of the overall volume. Forming the final 2D image by summing across planes requires synchronization across the GPUs. Iteration times include the forward simulation of the PSF, simulating the 2D image by convolving the PSF with the sample, reconstructing the 3D volume using the neural network, and jointly updating the parameters of both the neural network and the phase mask using the rectified Adam optimizer.

### Computer generated holography using deep learning (Figure 3f-l)

#### Visualized results

All procedures and data follow the original work by Eybposh et al.^12^ We generate a set of target patterns of 2048 × 2048 pixels for which we want to generate holograms. Each target pattern consists of three planes of dots with varying intensities. The goal is to generate a hologram (phase mask pattern) that will reproduce these three target patterns at the three desired image planes. The system we model for holography consists of a phase mask (quantized to 8 bits) illuminated by a plane wave, followed by a Fourier transforming lens to bring the generated pattern to the first image plane^12^. We simulate the resulting pattern for these three planes at -10, 0, or 10 mm from the focal plane of the lens.

We train a neural network to generate these holograms by predicting the complex field (amplitude and phase) at the first image plane given the target pattern^12^. We then propagate this field back to the phase mask plane in order to obtain the phase values for the pixels of the phase mask^12^. This phase mask is propagated to the three image planes to obtain the resulting pattern^12^. The network parameters are updated according to the accuracy loss between the resulting pattern and the target pattern, as in Eybposh et al.^12^ This training is performed in batches of 16 patterns at a time.

#### Scalability results

We report iteration times for training on a single batch of 16 patterns. Iteration time measures the time to perform a forward pass of the neural network, propagate the result to the phase mask plane to compute the phase mask pixel values, propagate to the three target image planes, and finally update the parameters of the neural network based on the accuracy loss between the simulated patterns and the target patterns. Parallelization is performed by splitting up the batch of 16 patterns across 1, 2, 4, or 8 GPUs (NVIDIA H100s).

### Multicolor PSF engineering (Figure 4a-f)

We demonstrate how a single phase mask can be engineered for spectral imaging of 25 different wavelengths ranging from 400 to 650 nm using a single-channel camera image. We simulate a programmable widefield fluorescence microscope and optimize its PSF to enable reconstruction of point sources emitting light of different wavelengths per point (at the focal plane). The reconstruction is performed using a FourierNet neural network ^3^, which involves a global convolution (implemented as a Fourier convolution for efficiency) followed by direct convolution layers with smaller filters. We jointly train both the reconstruction network and the pixels of the phase mask. The reconstruction network is able to exploit the sparsity of the sample as well as the PSF optimized to encode these specific wavelengths via fringe patterns in order to accurately reconstruct the points at their respective wavelengths.

### Computer generated holography through scattering media (Figure 4g-m)

We demonstrate the optimization of a hologram through scattering tissue rather than a homogeneous medium, as is typically assumed in computer generated holography. This simulation follows a typical Fourier holography system, similar to the system used in our demonstration of computer generated holography using deep learning. However, here we add the multislice scattering sample model that was used for refractive index microscopy and we also optimize the phase mask directly via gradient descent based on the Pearson correlation loss between the simulated pattern and the target pattern. Here, we are not relaying a focal plane to the camera but rather directly computing the intensity of the resulting pattern at each plane. The target pattern here is a 3D distribution of points of uniform intensity. For the scattering tissue, we use a volume 50 × 1200 × 1200 voxels large containing a reconstruction of a *C. elegans head*. We artificially scale the refractive index values in the sample by a factor of 5x in order to highlight the scattering for this demonstration, even though this is not a realistic sample. We first optimize a hologram through free space, using angular spectrum propagation through the same distance as the thickness of the scattering volume. We then simulate this hologram through both free space and through the scattering volume using the multislice beam propagation method. Finally, we optimize a hologram through the scattering volume and again simulate this hologram through the scattering volume to demonstrate that the intensity of the resulting points remains uniform throughout the volume, whereas the hologram optimized using the typical free space model results in degradation of the resulting pattern.

### Scalability results (Figure 5)

For ring deconvolution microscopy, computational aberration correction, refractive index microscopy, snapshot PSF engineering, and computer generated holography we present a comparison of timing between the original implementations and the Chromatix implementations. Generally, we are reporting the average time to perform one iteration of training or optimization when solving these inverse problems. We show these results in Figure 5 as relative performance versus number of GPUs used to parallelize each method. For all methods, outlier iteration times (3 standard deviations higher than the mean) were discarded prior to plotting which can lead to different numbers of samples for each point shown in Figure 5. Different methods are also simulated on different numbers of GPUs in parallel depending on how many GPUs could cleanly parallelize the workload for a given problem. We collect the following numbers of iteration times for the various methods: snapshot PSF engineering 1 GPU: 499 iterations, 2 GPUs: 499 iterations, 4 GPUs: 499 iterations, 8 GPUs: 499 iterations; ring deconvolution 1 GPU: 4980 iterations, 2 GPUs: 4983 iterations, 4 GPUs: 4981 iterations, 8 GPUs: 4992 iterations; refractive index microscopy 1 GPU: 24919 iterations, 2 GPUs: 12530 iterations, 3 GPUs: 8362 iterations, 4 GPUs: 6284 iterations, 6 GPUs: 4192 iterations, 7 GPUs: 3494 iterations, 8 GPUs: 3128 iterations; computer generated holography 1 GPU: 99 iterations, 2 GPUs: 100 iterations, 4 GPUs: 99 iterations, 8 GPUs: 98 iterations; computational aberration correction 1

GPU: 998 iterations, 2 GPUs: 995 iterations, 3 GPUs: 996 iterations, 4 GPUs: 996 iterations, 5 GPUs: 991 iterations, 6 GPUs: 996 iterations, 7 GPUs: 992 iterations, 8 GPUs: 998 iterations. Note that refractive index microscopy performs fewer iterations scaling inversely with number of GPUs available due to greater numbers of GPUs supporting larger batch sizes to be optimized at a time. See the corresponding sections in the methods for more details on the scalability results for each application.

## Data Availability

We use publicly available data from prior work for all experiments shown. For ring deconvolution (Figure 2a-f)^52^, the data we use can be obtained from: https://github.com/apsk14/rdmpy. For computational aberration correction (Figure 2g-n)^47^, the data we use can be obtained from: https://github.com/iksungk/CoCoA. The measured wavefront for computational aberration correction (Figure 2l) was obtained upon request from Kang et al.^47^ For refractive index microscopy (Figure 2o-u)^17^, the data we use can be obtained from: https://dataverse.tdl.org/dataverse/DMD-MLA-01. For snapshot PSF engineering (Figure 3a-e)^3^, the data we use can be obtained from: https://doi.org/10.25378/janelia.25277269.v1. For holography through scattering (Figure 4g-m), the data can be obtained from https://github.com/Waller-Lab/multislice^16^. Any other data shown was completely programmatically generated in simulation.

## Code Availability

The Chromatix library is available at https://github.com/chromatix-team/chromatix. The documentation and installation instructions are available at https://chromatix.readthedocs.io. This work used version 0.4, and Chromatix is distributed under the MIT License.

## Extended Data

### Comparison of features against other simulators

**Table 1.**
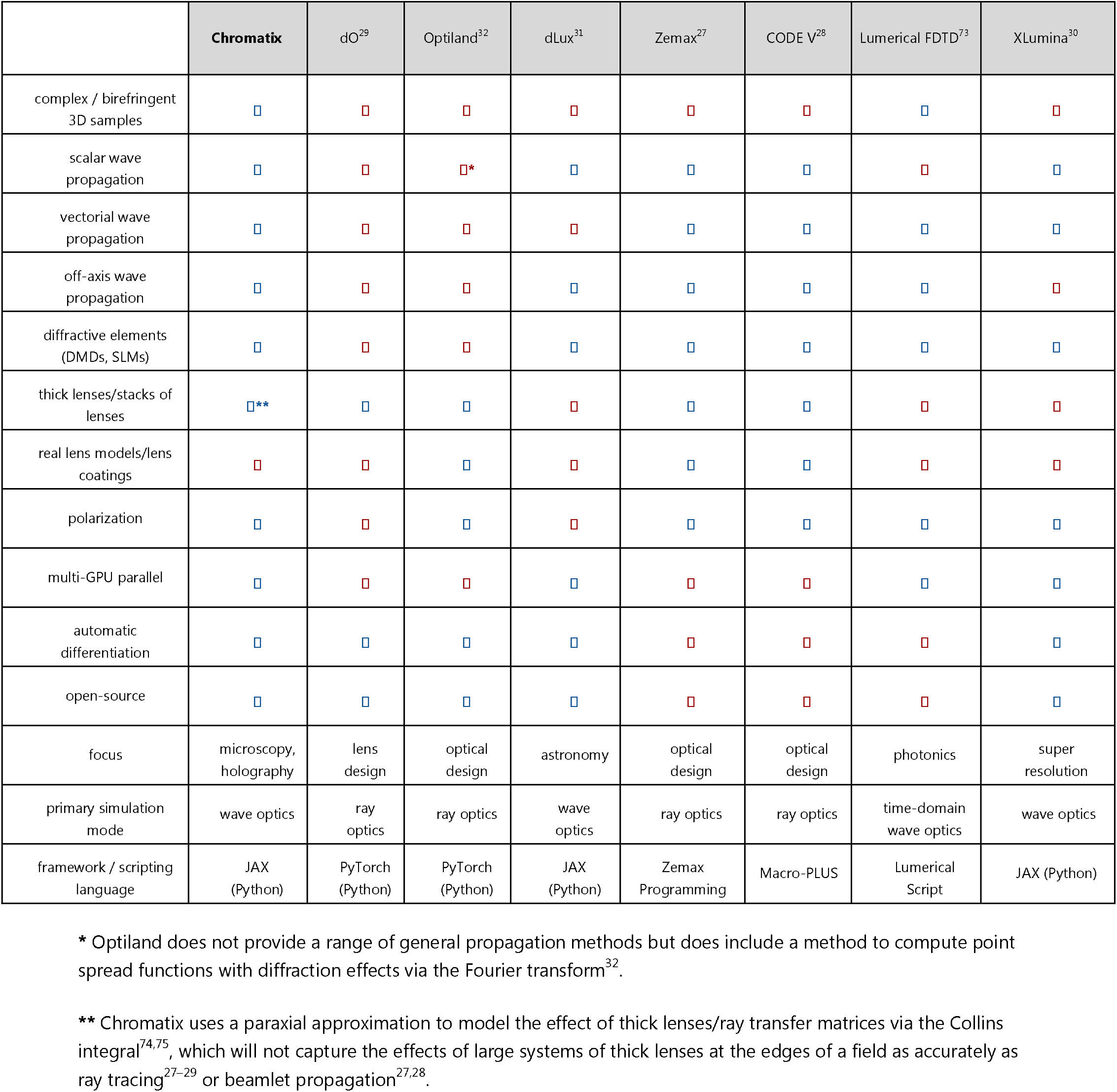
Comparison of features against other simulation libraries/software.

## Extended figures

**Fig. 1.**
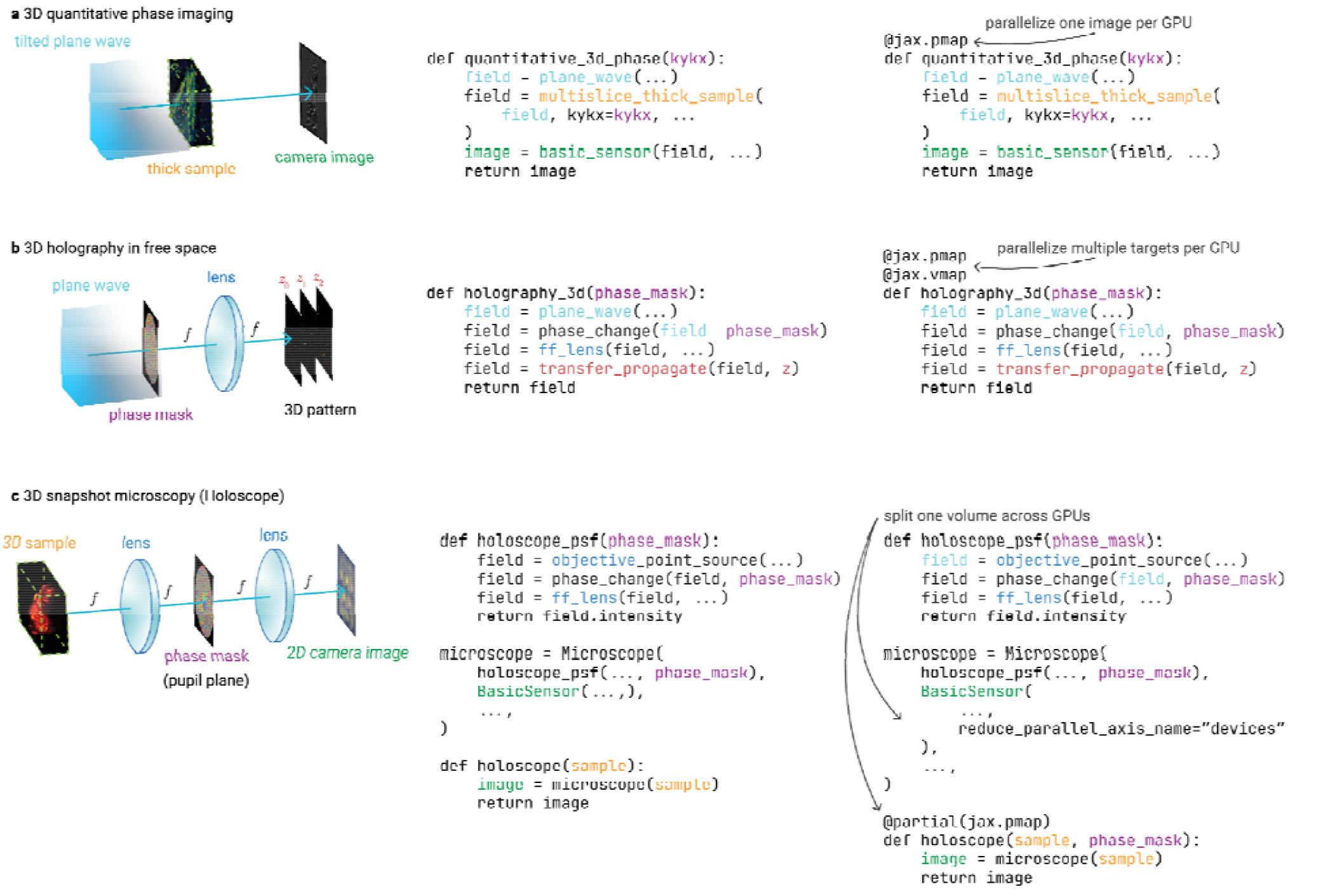
Chromatix enables parallelization of optical simulations with minimal code changes. For all methods, we show the optical model followed by its implementation using Chromatix in both an unparallelized single GPU simulation as well as the parallelized versions that run across multiple GPUs. This display demonstrates how Chromatix allows a researcher to describe an optical model independently of the code required to scale that model across multiple GPUs in parallel. Colors in the code denote the corresponding elements in the model diagrams on the left. Note that the code shown is not the complete code that is actually used in demonstrations across the rest of the paper; this code is meant to provide a high level depiction of Chromatix code and parallelization. Ellipses indicate code that has been intentionally omitted for brevity. Simulation results are identical for all of the parallelization scenarios demonstrated here. **a**, Optical model for quantitative phase imaging showing how a single simulation can be easily run independently on multiple GPUs. **b**, Optical model for 3D holography showing how a simulation can be run multiple times in parallel on one GPU, with this whole batch of simulations run in parallel across multiple GPUs. **c**, Optical model for 3D snapshot microscopy (the Holoscope) showing how a simulation can be run across one large 3D sample split across multiple GPUs to enable larger simulations/optimizations.

**Fig. 2.**
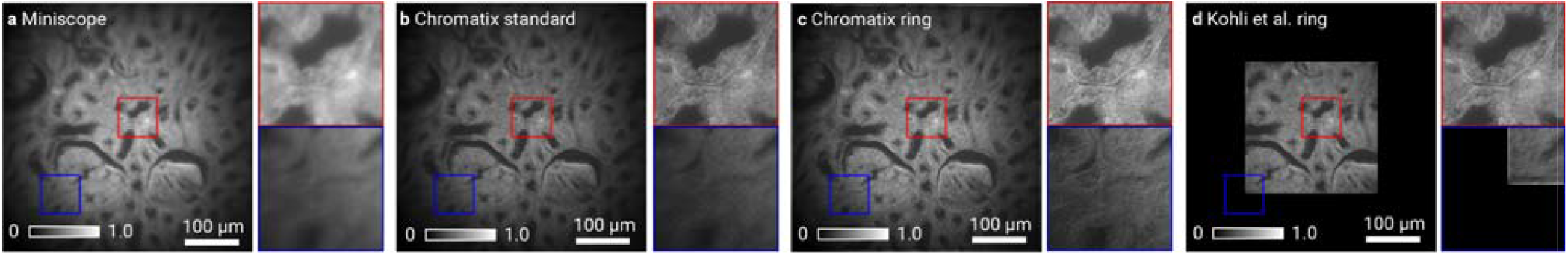
Reconstruction performance of ring deconvolution without correction of vignetting. Highlighted regions in **a** - **d** show zoomed in cutouts from the center (red) and edge (blue) of the field of view on the bottom row. Intensity values in **a** - **d** are normalized. Measured image of incoherently illuminated rabbit liver **a** from a Miniscope with a GRIN lens showing extreme spatially-varying aberrations across the field of view; **b**, Chromatix rotationally invariant deconvolution demonstrating improved image quality across the entire field of view; **c**, Spatially invariant (standard) deconvolution demonstrating good quality in the center but degraded quality in the edges of the field of view; and **d**, original PyTorch rotationally invariant deconvolution which does not parallelize and cannot fit the whole field of view on 1 H100 GPU (80 GB memory).

**Fig. 3.**
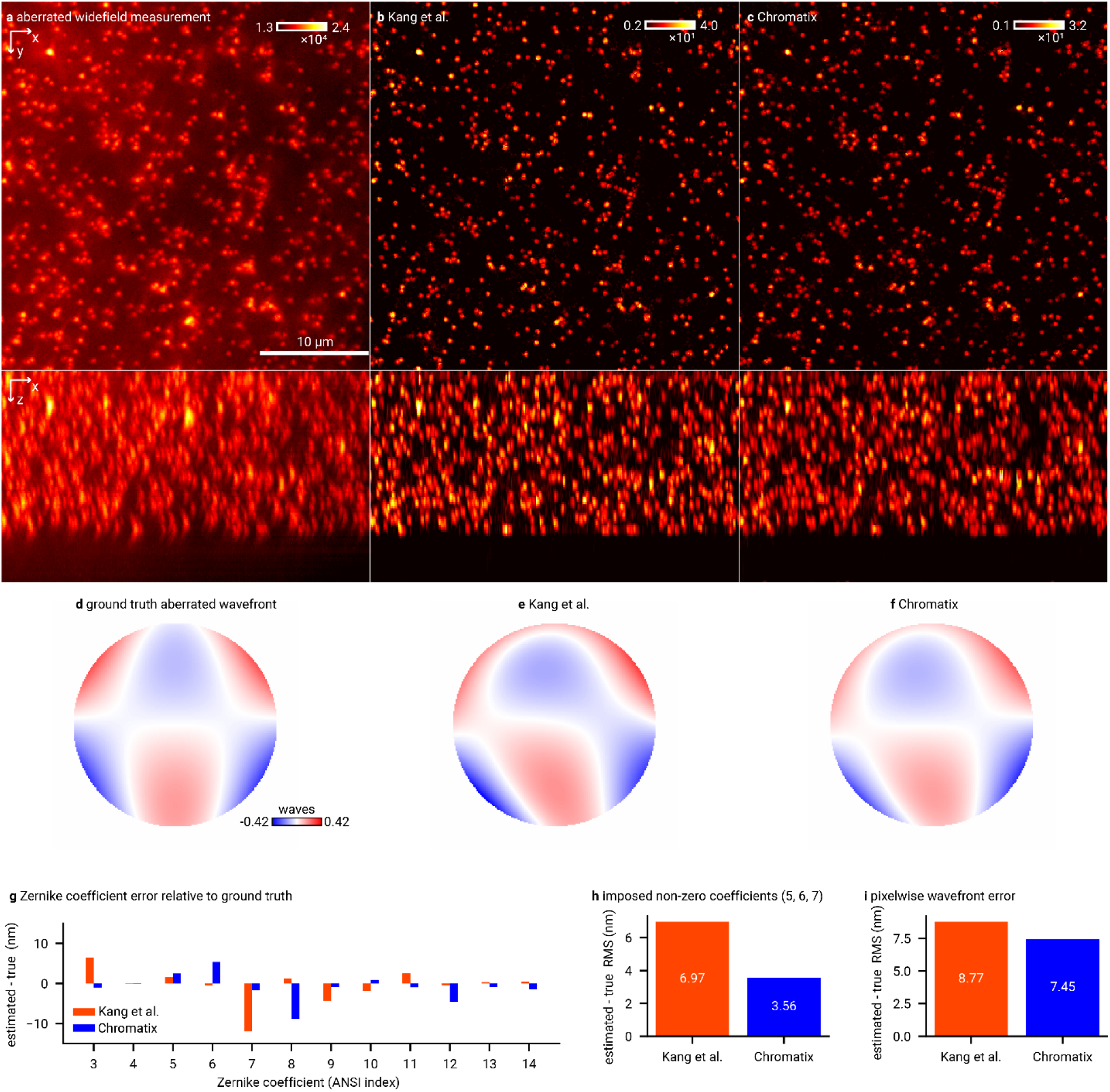
Chromatix outperforms the original CoCOA implementation for estimating aberrations on fluorescent bead data. **a**, Widefield measurement of fluorescent beads, showing aberrations and high background. **b**, Kang et al.^47^ reconstruction of the fluorescent beads. **c**, Chromatix reconstruction of the fluorescent beads. **d**, Ground truth aberrated wavefront based on the imposed Zernike modes. **e**, Kang et al.^47^ estimation of the aberrated wavefront. **f**, Chromatix estimation of the aberrated wavefront. **g**, Bar plot comparing the wavefront error (nm) between the true imposed coefficients and the estimated coefficients for each Zernike mode across both Chromatix (blue) and the original implementation^47^ (red). Note that all Zernike modes aside from 5, 6, and 7 should have a coefficient of 0 nm in the imposed aberration. **h**, Bar plot of root mean square (RMS) of the difference between the true wavefront coefficients and the estimated coefficients across the three non-zero Zernike coefficients of the imposed aberration, demonstrating that Chromatix has approximately half the error compared to the Kang et al. implementation^47^. **i**, Bar plot of RMS of the pixelwise difference between the estimated aberrated wavefront and the true imposed wavefront, showing the wavefront recovered by Chromatix is more accurate to the ground truth than the original implementation^47^.

**Fig. 4.**
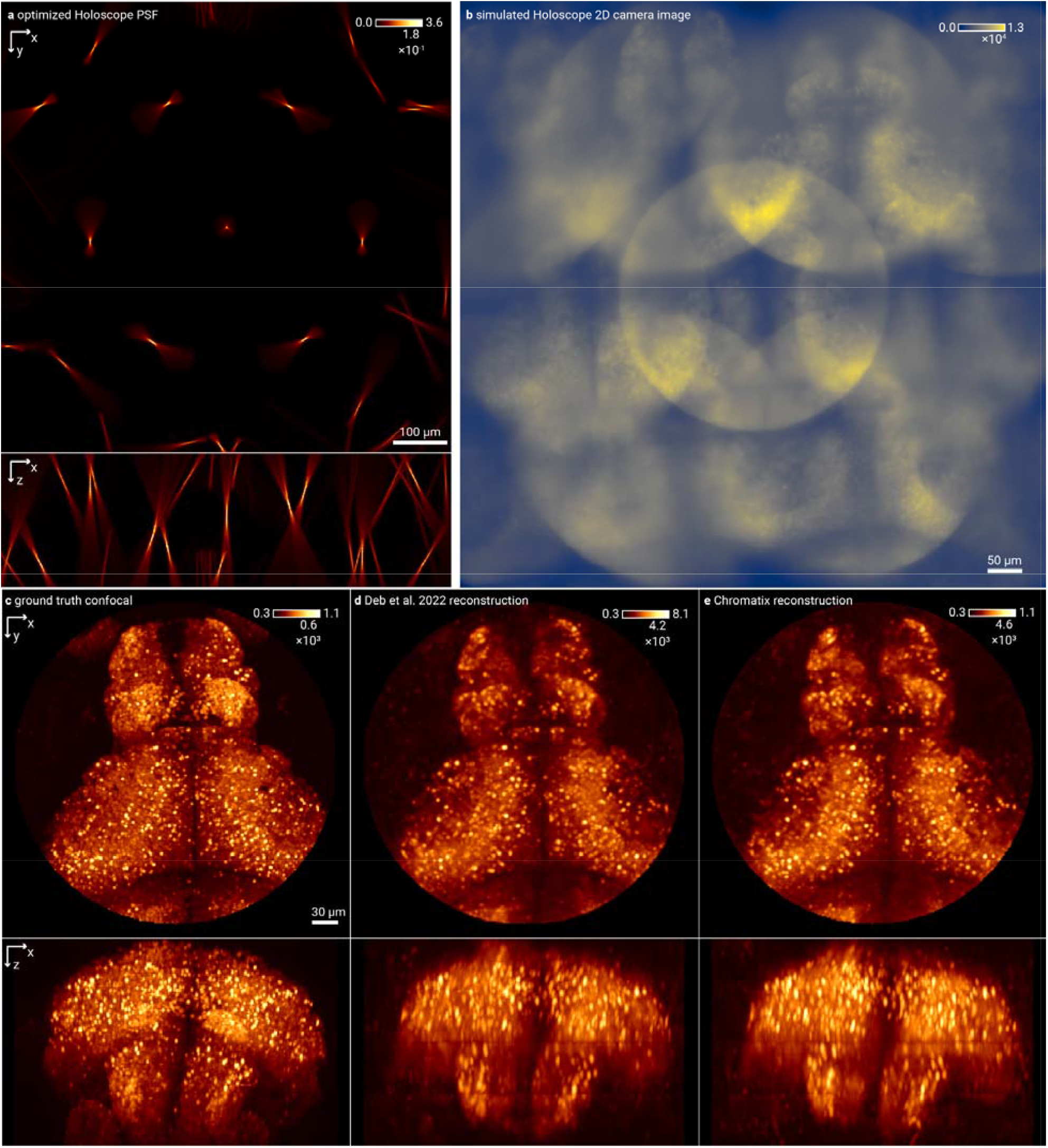
Chromatix matches the *in silico* Holoscope reconstruction performance of Deb et al. **a**, Holoscope PSF optimized for zebrafish with a circular aperture in the original implementation by Deb et al.^3^ **b**, Simulated camera image showing a 3D zebrafish sample captured in a single 2D image. **c**, Ground truth confocal volume of the brain of a larval zebrafish (*D. rerio*), which will be over an order of magnitude slower to capture than the single snapshot image in (**b**). **d**, Reconstruction using the original implementation by Deb et al.^3^ **e**, Reconstruction using Chromatix, which matches the reconstruction performance of the original implementation despite being significantly faster computationally as shown in Figure 5. On a test set of 10 volumes, reconstruction networks trained with identical PSFs offer an SSIM of 0.979 ± 0.003 (not significantly different at p = 0.695 via two-sided t-test) for both Chromatix and the original implementation^3^ and a peak signal to noise ratio (PSNR) of 35.91 ± 0.92 for Chromatix versus 36.99 ± 1.07 for the original implementation^3^ (not significantly different at p = 0.063 via two-sided t-test). For both SSIM and PSNR, higher is better.

**Fig. 5.**
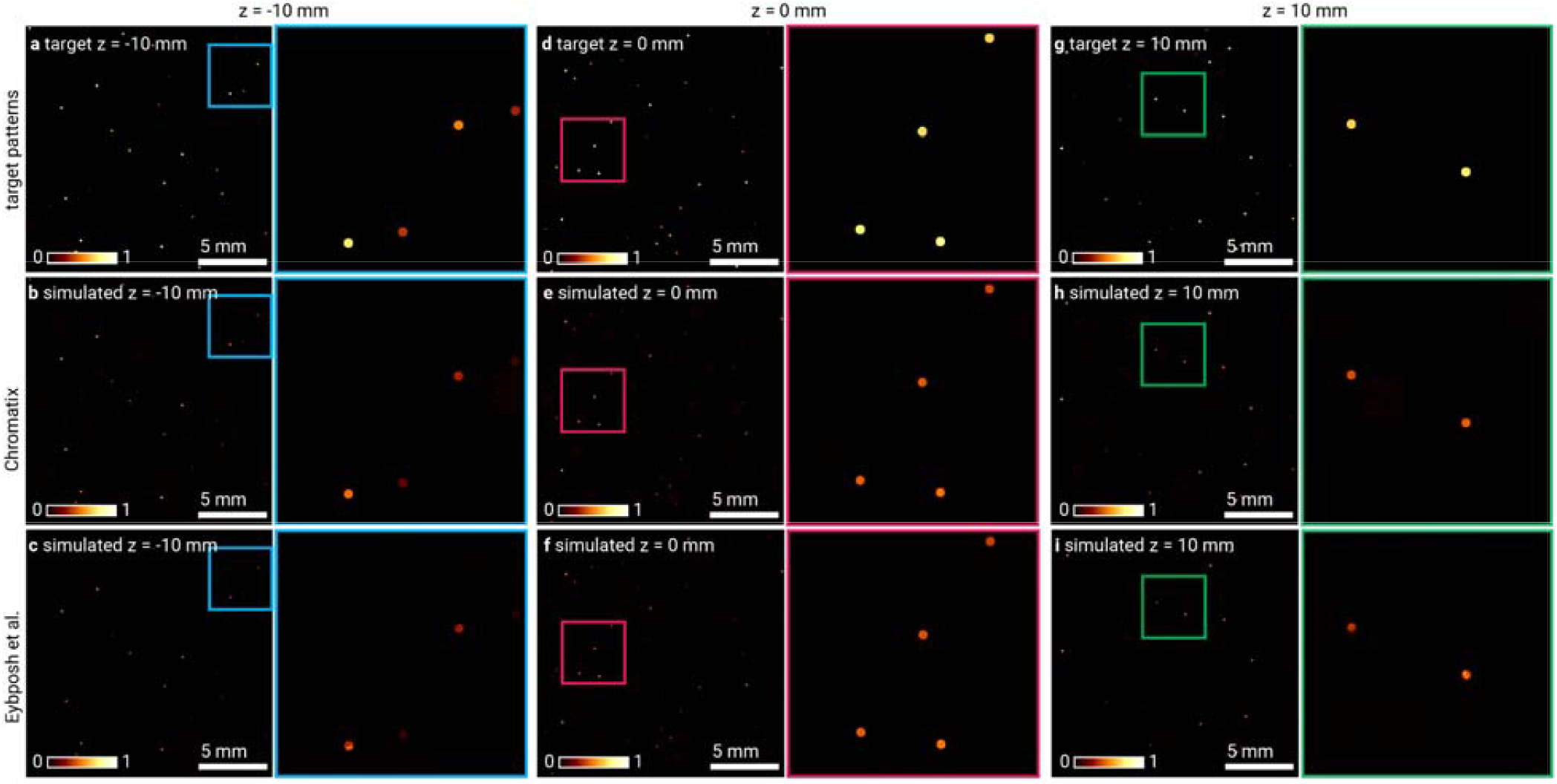
Chromatix matches the *in silico* DeepCGH performance of Eybposh et al. **a**,**d**,**g**, Target pattern of dots of random intensity at planes -10 mm, 0 mm, and 10 mm from the focal plane respectively. **b**,**e**,**h**, Simulated pattern using holograms generated by a DeepCGH^12^ model trained using Chromatix at planes -10 mm, 0 mm, and 10 mm from the focal plane respectively. **c**,**f**,**i**, Simulated pattern using holograms generated by a DeepCGH model trained using the original implementation by Eybposh et al.^12^ at planes -10 mm, 0 mm, and 10 mm from the focal plane respectively. On a test set of 16 target patterns, we obtain an SSIM of 0.985 ± 0.001 for Chromatix versus 0.982 ± 0.001 for the original implementation^12^ (significantly different at p = 0.018 < 0.05 via two-sided t-test) and a PSNR of 35.40 ± 0.37 for Chromatix versus 34.95 ± 0.16 for the original implementation^12^ (not significantly different at p = 0.177 via two-sided t-test). For both SSIM and PSNR, higher is better.

## Notes

https://chromatix.readthedocs.io/en/latest/

https://github.com/chromatix-team/chromatix

### Competing Interest Statement

The authors have declared no competing interest.

### Summary of Updates

This version of the manuscript has been updated with reduced word counts and a condensed Figure 2. Other minor formatting changes have been made in preparation for publication.

