## Extended Data Figure 1 for "Chromatix: a differentiable, GPU-accelerated wave-optics library"

**a** 3D quantitative phase imaging

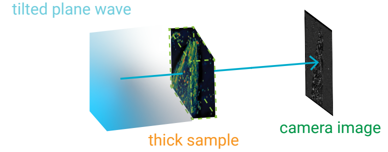


```
def quantitative_3d_phase(kykx):
    field = plane_wave(...)
    field = multislice_thick_sample(
        field, kykx=kykx, ...
    )
    image = basic_sensor(field, ...)
    return image
```

parallelize one image per GPU

```
@jax.pmap
def quantitative_3d_phase(kykx):
    field = plane_wave(...)
    field = multislice_thick_sample(
        field, kykx=kykx, ...
    )
    image = basic_sensor(field, ...)
    return image
```

**b** 3D holography in free space

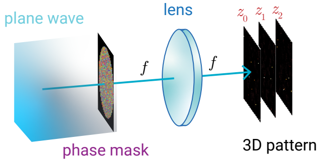


```
def holography_3d(phase_mask):
    field = plane_wave(...)
    field = phase_change(field  phase_mask)
    field = ff_lens(field, ...)
    field = transfer_propagate(field, z)
    return field
```

parallelize multiple targets per GPU

```
@jax.pmap
@jax.vmap
def holography_3d(phase_mask):
    field = plane_wave(...)
    field = phase_change(field, phase_mask)
    field = ff_lens(field, ...)
    field = transfer_propagate(field, z)
    return field
```

**c** 3D snapshot microscopy (Holoscope)

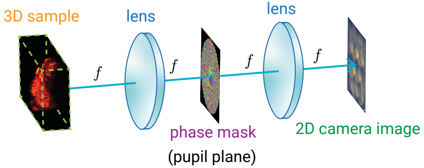


```
def holoscope_psf(phase_mask):
    field = objective_point_source(...)
    field = phase_change(field, phase_mask)
    field = ff_lens(field, ...)
    return field.intensity

microscope = Microscope(
    holoscope_psf(..., phase_mask),
    BasicSensor(...,),
    ...,
)

def holoscope(sample):
    image = microscope(sample)
    return image
```

split one volume across GPUs

```
def holoscope_psf(phase_mask):
    field = objective_point_source(...)
    field = phase_change(field, phase_mask)
    field = ff_lens(field, ...)
    return field.intensity

microscope = Microscope(
    holoscope_psf(..., phase_mask),
    BasicSensor(
        ...,
        reduce_parallel_axis_name="devices"
    ),
    ...,
)

@partial(jax.pmap)
def holoscope(sample, phase_mask):
    image = microscope(sample)
    return image
```
