## Supplementary figures and images for "Chromatix: a differentiable, GPU-accelerated wave-optics library"

### Extended Data Figure 2

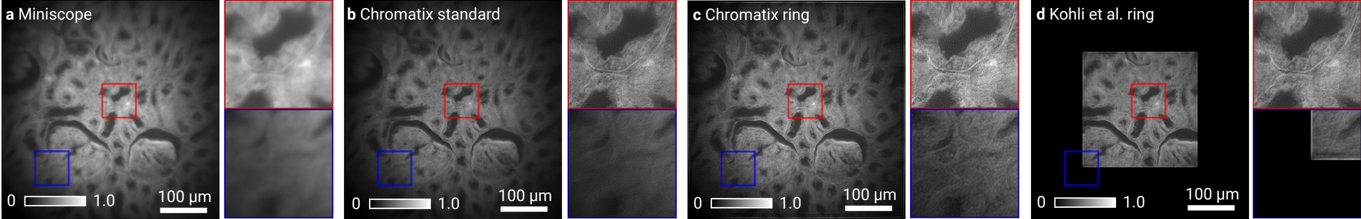

### Extended Data Figure 3

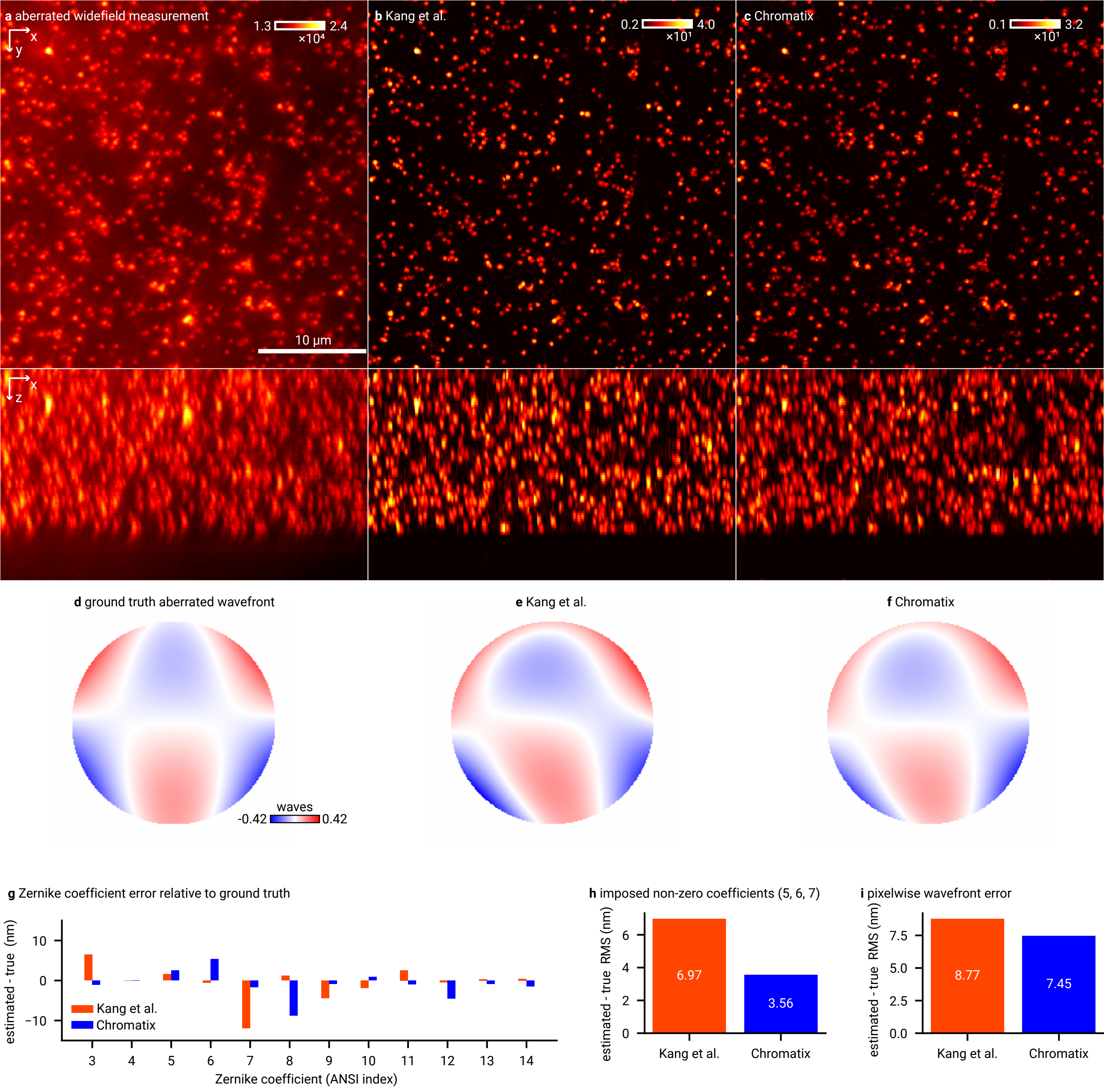

### Extended Data Figure 4

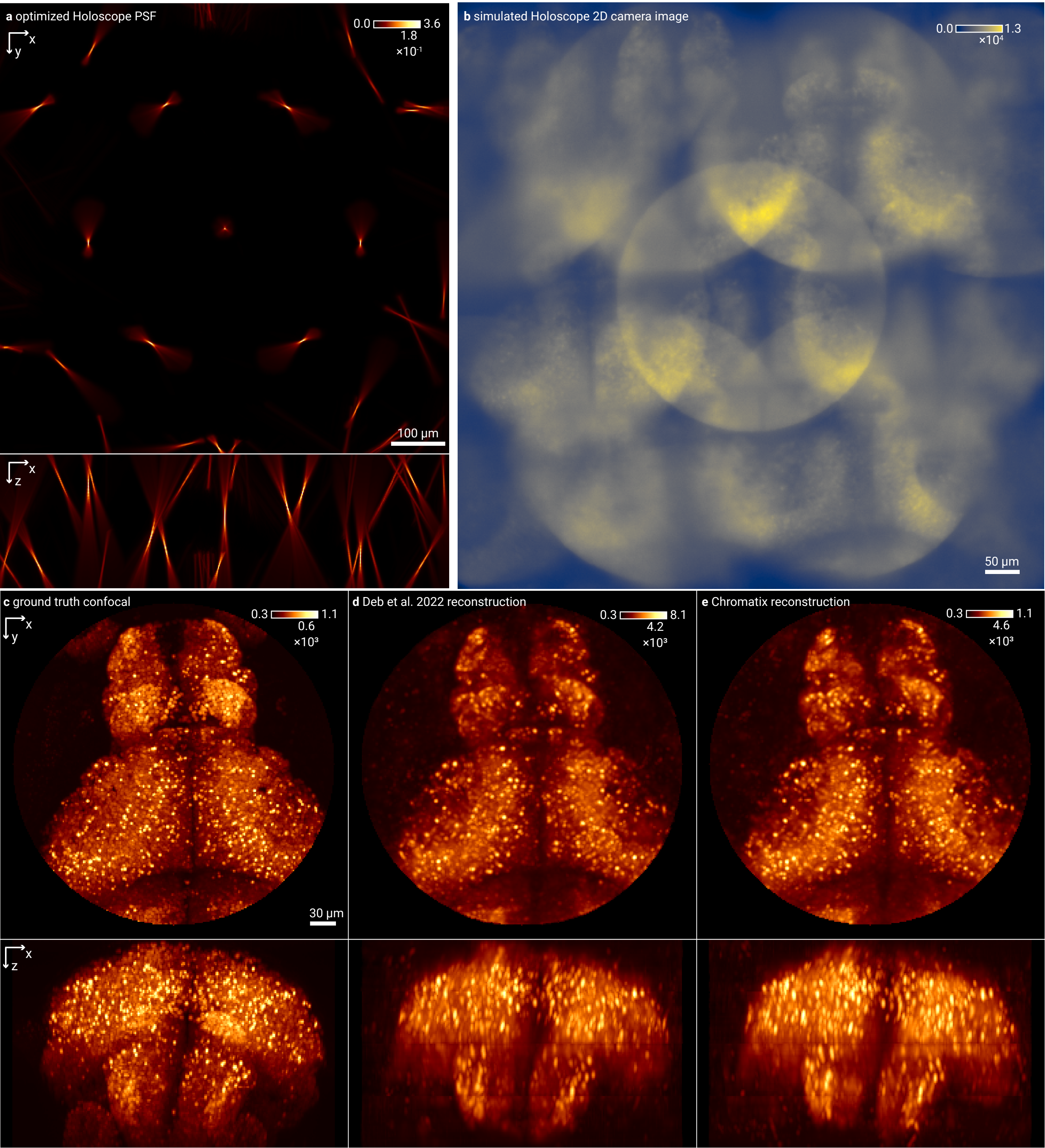

### Extended Data Figure 5

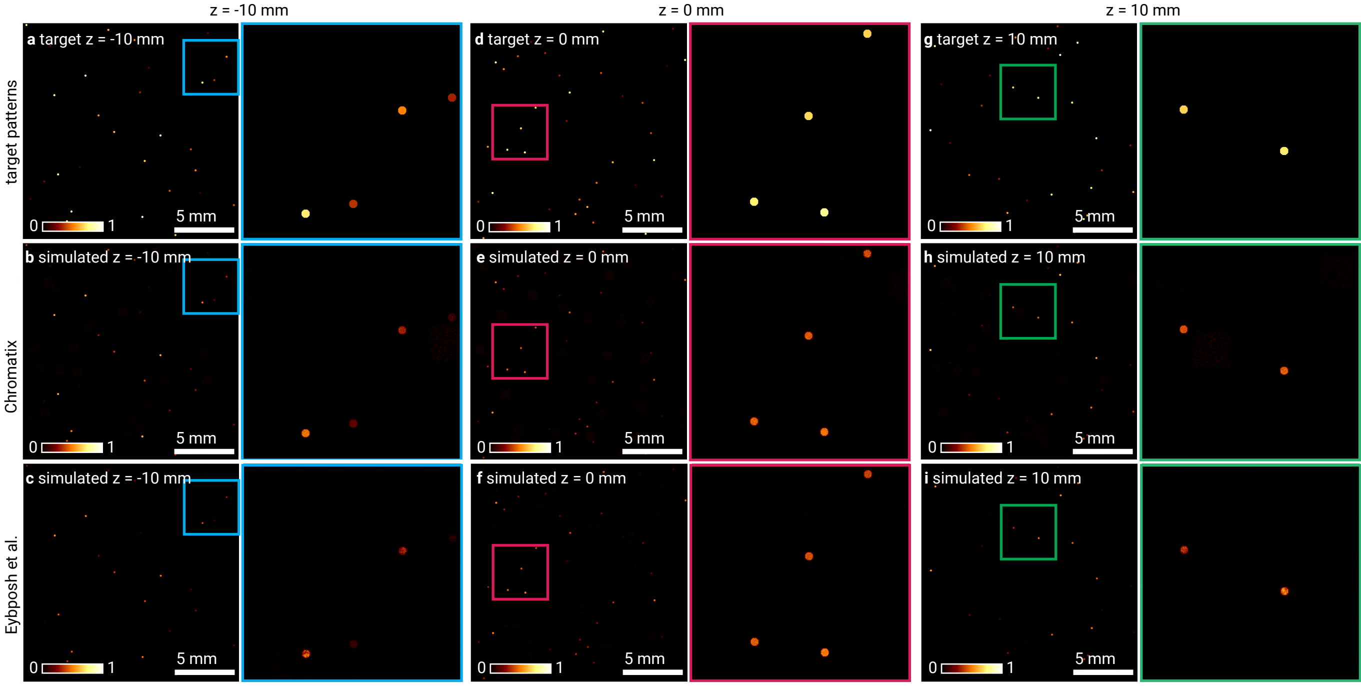
