## Extended Data Table 1 for "Chromatix: a differentiable, GPU-accelerated wave-optics library"

**Table 1 | Comparison of features against other simulation libraries/software.**

| | **Chromatix** | dO[29] | Optiland[32] | dLux[31] | Zemax[27] | CODE V[28] | Lumerical FDTD[73] | XLumina[30] |
|---|---|---|---|---|---|---|---|---|
| complex / birefringent 3D samples | ✓ | □ | □ | □ | □ | □ | ✓ | □ |
| scalar wave propagation | ✓ | □ | □* | ✓ | ✓ | ✓ | □ | ✓ |
| vectorial wave propagation | ✓ | □ | □ | □ | ✓ | ✓ | ✓ | ✓ |
| off-axis wave propagation | ✓ | □ | □ | ✓ | ✓ | ✓ | ✓ | □ |
| diffractive elements (DMDs, SLMs) | ✓ | □ | □ | ✓ | ✓ | ✓ | ✓ | ✓ |
| thick lenses/stacks of lenses | ✓** | ✓ | ✓ | □ | ✓ | ✓ | □ | □ |
| real lens models/lens coatings | □ | □ | ✓ | □ | ✓ | ✓ | □ | □ |
| polarization | ✓ | □ | ✓ | □ | ✓ | ✓ | ✓ | ✓ |
| multi-GPU parallel | ✓ | □ | □ | ✓ | □ | □ | ✓ | ✓ |
| automatic differentiation | ✓ | ✓ | ✓ | ✓ | □ | □ | □ | ✓ |
| open-source | ✓ | ✓ | ✓ | ✓ | □ | □ | □ | ✓ |
| focus | microscopy, holography | lens design | optical design | astronomy | optical design | optical design | photonics | super resolution |
| primary simulation mode | wave optics | ray optics | ray optics | wave optics | ray optics | ray optics | time-domain wave optics | wave optics |
| framework / scripting language | JAX (Python) | PyTorch (Python) | PyTorch (Python) | JAX (Python) | Zemax Programming | Macro-PLUS | Lumerical Script | JAX (Python) |

***** Optiland does not provide a range of general propagation methods but does include a method to compute point spread functions with diffraction effects via the Fourier transform[32].

****** Chromatix uses a paraxial approximation to model the effect of thick lenses/ray transfer matrices via the Collins integral[74,75], which will not capture the effects of large systems of thick lenses at the edges of a field as accurately as ray tracing[27–29] or beamlet propagation[27,28].
